# Critical phenomenon underlies *de novo* luminogenesis during mammalian follicle development

**DOI:** 10.1101/2025.09.09.674793

**Authors:** Kim Whye Leong, Yuting Lou, Arikta Biswas, Jue Yu Kelly Tan, Boon Heng Ng, Xixun Lu, Xin Ping Joan Teo, Thong Beng Lu, Carlo Bevilacqua, Isabelle Bonne, Robert Prevedel, Tetsuya Hiraiwa, Chii Jou Chan

## Abstract

A key process during mammalian folliculogenesis is the formation of a fluid-filled antrum around the oocyte. While it is known that the antrum is critical for oocyte maturation and the eventual ovulation, the detailed process underlying *de novo* luminogenesis remains poorly understood. In this study, we investigated the spatiotemporal dynamics of lumen growth and the cellular mechanism driving this process, using advanced microscopy, molecular perturbations and computational modelling. We found that in secondary follicles, the interstitial gaps exist in a near-critical regime, characterised by a highly dynamic and interconnected fluid network with gap size obeying a power-law distribution. Above a critical size of 180 μm, we observed an onset of cell death and the emergence of a stably growing dominant fluid cavity resembling phase separation. By modelling the granulosa cells and the interstitial fluid as a binary fluid, we reproduced the near-critical and phase-separated regimes and found that a spatial gradient of cell-fluid interfacial tension, as observed experimentally, is sufficient to robustly maintain the secondary follicles in a near-critical state. Reducing cell-fluid interfacial tension globally by weakening the cell-cell adhesion between granulosa cells leads to fragmentation of fluid and growth arrest of the dominant cavity, as predicted by the model. Importantly, perturbing the critical state in secondary follicles leads to impaired follicle growth. Altogether, our study reveals how collective spatiotemporal regulation of cell junction mechanics and cell death can contribute to the effective transition between distinct regimes of fluids, which are indispensable for functional maturation of follicles.

## MAIN TEXTS

The development of functional oocytes within follicles, or folliculogenesis, is a critical process in mammalian reproduction^1,2^. Such process is regulated by various biochemical and biophysical signals in the follicular microenvironment^1,3^. Starting from the secondary follicle stage, the oocyte is surrounded by multi-layered granulosa cells (GC) and an outer layer of contractile theca cells^1^. As the follicle continues to grow, fluid-filled cavities (lumen) appear and eventually merge to become a large cavity known as antrum. While such process is conserved across mammalian species, the antrum size varies from species to species, with that in the larger species, such as human and bovine, occupying more than 95% of the follicle during ovulation^4^, while the follicles in the small mammals, such as mice, contain less follicular fluid^5^.

Despite previous correlational studies linking antrum formation to oocyte maturation^6,7^, and speculation that the follicular fluid may mediate paracrine signalling between the oocyte and GCs^6,8^the biological functions of follicular lumen remain enigmatic. Furthermore, the spatiotemporal dynamics of lumen formation is poorly characterised, potentially due to the lack of approaches to study these events from a collective perspective. While past studies have posited that cell death may regulate lumen expansion through osmotic means^5,9^, the exact molecular mechanisms driving *de novo* lumenogenesis remain unclear. In this study, we examined early phases of lumen formation in secondary and tertiary follicle stages, and the functional impact of lumen in follicle growth. By combining quantitative timelapse imaging, molecular perturbations and biophysical modelling, our study provides a new framework to understand lumen formation in folliculogenesis through the lens of critical phenomena.

### Interstitial gaps grow during secondary-tertiary follicle development

To better understand the temporal events of luminogenesis in murine follicles, we first performed immunofluorescence staining on ovarian tissue slices to identify lumenal structures in follicles (Fig. 1a). In follicles with size ranging between 100 and 180 μm, which we term ‘secondary follicles’, we observed actin-labelled canal-like gaps between granulosa cells (GCs). However, in follicles larger than 180 μm, which we termed ‘tertiary follicles’, such interconnected gaps are less visible. To ascertain that these structures are true fluid gaps, we cultured, *in vitro*, intact follicles isolated from ovaries and directly visualised the interstitial gaps (IGs) using a dextran-FITC based fluorescence exclusion method routinely used in the study of lumen^10,11^. With this method, we discovered an extensive network of IGs in follicles of all sizes. Apart from the distinct lumen described in Fig. 1a, we observed smaller IGs that were previously inconspicuous in actin-stained tissue sections. In addition, we found that FITC also resides in blob-like structures with a size comparable to that of a GC (Fig. 1b). We hypothesised that these structures were dead GCs with compromised cell membrane, allowing FITC to fill the entire cell body. Using the positive FITC signals, we were able to exclude the blob-like structures and segment the IGs within the entire follicle, using Imaris and a custom Python pipeline (see Methods, Extended Data Fig. 1a, b; Supplementary Video 1). With three-dimensional rendering, we found a positive correlation between IG number and IG fraction with follicle diameter, suggesting that the IGs are being actively generated during follicle growth (Fig. 1c). We next quantified the largest gap volume as a function of follicle size and found that the size of IGs varies across multiple length scales, with the largest IG volume typically 30-40 times larger than the mean IG volume (Fig. 1d). Of note, the largest IG volume growth scales strongly with the follicle size, suggesting that the largest IG dominates overall fluid expansion, and will likely become the dominant cavity in the later stage of antrum formation.

**Figure 1:**
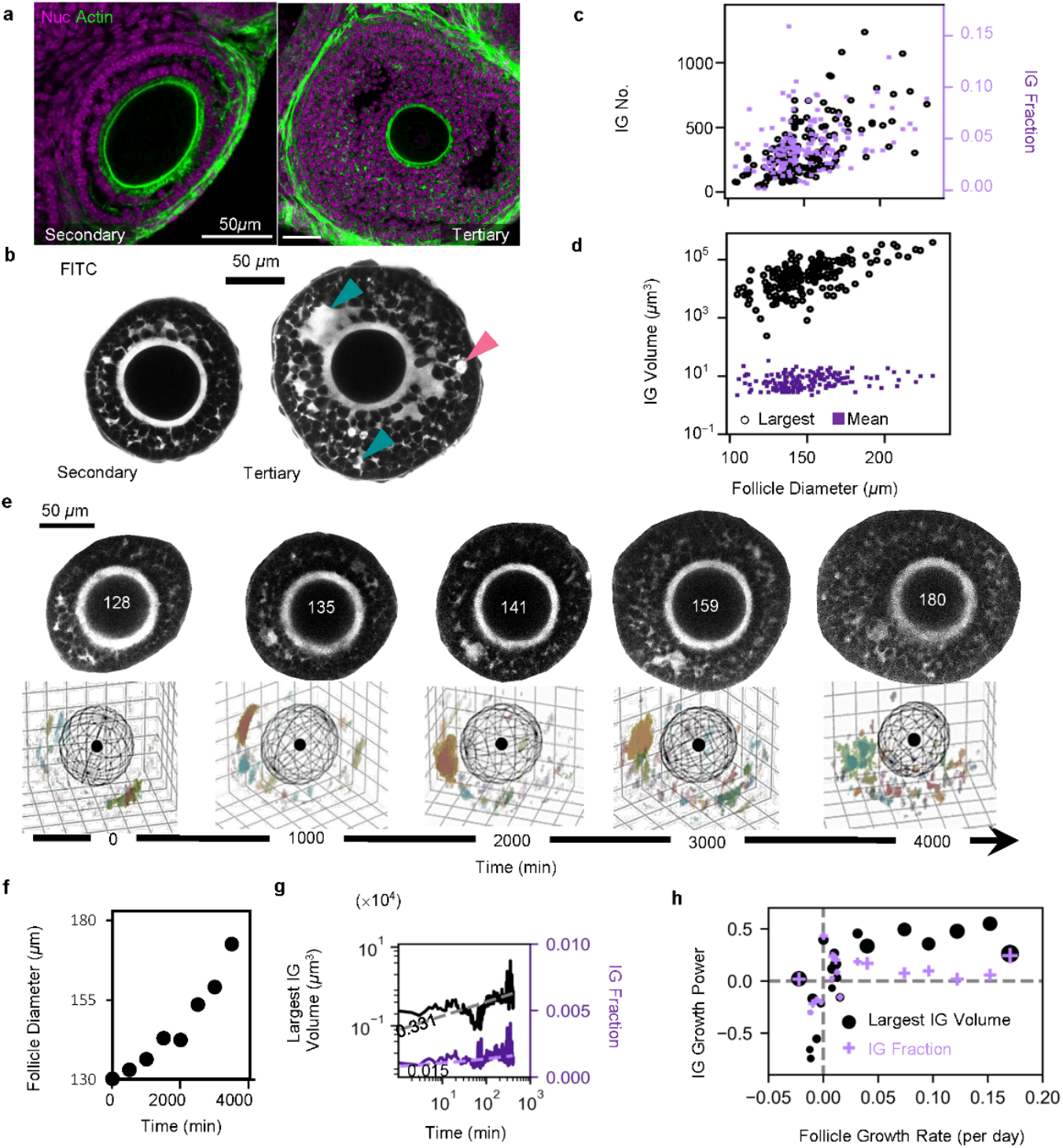
Growth of interstitial gaps (IGs) correlates with follicle growth during secondary-tertiary follicle development. **a, b**, Representative images of secondary (left, 129 µm (top), 139 µm (bottom)) and tertiary (right, 300 µm (top), 190 µm (bottom)) follicles. Scale bar: 50 µm. **a**, Tissue slices stained with DAPI (nucleus, magenta) and phalloidin (actin, green). **b**, Isolated follicles stained with FITC. Green arrowheads represent IGs while pink arrowheads represent ‘FITC-blobs’. **c, d**, Population analysis of IG evolution on isolated follicles stained with FITC (N = 5, n = 153) **c**, Plot of total IG number (black circle, Pearson *R* = 0.646, *p* = 0) and total IG fraction (purple square, Pearson *R* = 0.420, *p* = 0) against follicle size. **d**, Plot of largest IG volume (black circle, Pearson *R* = 0.680, *p* = 0) and mean IG volume (purple square, Pearson *R* = 0.368, *p* = 0) against follicle diameter. **e**, Sequential images of a follicle stained with FITC (top) and the corresponding 3D-rendered IGs (bottom), tracked over 66 h. The numbers correspond to the follicle diameter at each time point. Each colour in the 3D-rendered image represents a distinct IG. See Supplementary Video 1 and 2. **f**, Plot of follicle diameter against time, for the corresponding follicle in **e. g**, Plot of the largest IG volume (black) and total IG fraction (purple) against time, for the corresponding follicle in **e**. Numbers correspond to the slope at the later phase of development. **h**, Plot of growth power of largest IG volume (black circle) and total IG fraction (purple cross) against mean follicle growth rate (N = 4, n = 21). Size of circles corresponds to the initial diameter of follicles.

To confirm our findings on IG temporal evolution based on population-based study, we performed long-term timelapse imaging to track the IG dynamics within a single follicle over several days (Fig. 1e, f, and Supplementary Video 2). We found, within 72 h of culture, a similar increase in the largest IG volume and IG fraction (Fig. 1g), corroborating results from our population-based study. To investigate the relationship between follicle and lumen growth, we plotted the fitted growth powers of both the largest IG volume and IG fraction against the follicle growth rate (See Methods, Fig. 1h). We found that lumen growth is linked to follicle growth, while lumen shrinkage, as indicated in negative growth powers of largest IG and IG fraction, is correlated with follicle growth arrest. Our results suggest that proper fluid expansion may be required for functional follicle growth. Importantly, we demonstrate the feasibility of our *in vitro* culture system for long-term study of luminogenesis during secondary-tertiary follicle development.

We next examined the IG fluid properties, using fluorescence recovery after photobleaching (FRAP). We found that the IGs and blobs have a recovery half-life of 1 and 1.5 s (Extended Data Fig. 2a), which correspond to a diffusion rate of about 1.5 μm^2^/s and 0.8 μm^2^/s, respectively (See Methods). These diffusion rates are on the same order of magnitude with that found from FRAP studies in glycerol solution^12^, suggesting that both the IGs and blobs are viscous fluids, and that the cell debris in blobs contribute to a higher viscosity. We further found that the recovery half-life of IGs is negatively correlated with follicle size (Extended Data Fig. 2b, c), suggesting that the IGs in larger follicles are less viscous than those in the smaller follicles. As a complementary approach to the low-frequency viscosity measurements in FRAP, we next applied Brillouin microscopy, an optoelastography technique known to probe tissue material properties^13,14^, to study IG micro-viscosity at high frequency. We found a significant decrease in Brillouin shift of IGs in tertiary follicles, with values similar to that of the surrounding medium (Extended Data Fig. 2d, e), suggesting that the IG viscosity resembles that of pure fluid. The slight decrease in the linewidth of IGs in tertiary follicles, despite being nominal (Extended Data Fig. 2e), also indicates lower micro-viscosity in these IGs. Together, our data indicates that the IGs are more viscous in IGs of secondary follicles.

### Luminogenesis is characterised by regime transition of IGs

We hypothesize that the distinct morphological features and fluid viscosity in the secondary and tertiary follicles reflect a fundamental difference in the collective dynamics of fluids in the two stages. To this end, we performed timelapse imaging with a higher spatiotemporal resolution to study the dynamical properties of IGs. We found that the IGs in the secondary follicles were highly dynamic compared to that of tertiary follicles (Fig. 2a, see Supplementary Video 3, 4). By quantifying the decay in autocorrelation of the FITC signals (Fig. 2b, see Methods), we extract the correlation time τ, which reflects the timescale at which the IGs retains its morphology. We found that the IGs in the secondary follicles have a shorter τ compared to that of tertiary follicles, indicating a more fluctuating nature of fluid in the secondary follicles. Indeed, τ increased with follicle size, eventually reaching a plateau value once the follicles grow beyond 200 μm (Fig. 2b, right), confirming that the IGs in secondary follicles are more dynamic than those in tertiary follicles.

**Figure 2:**
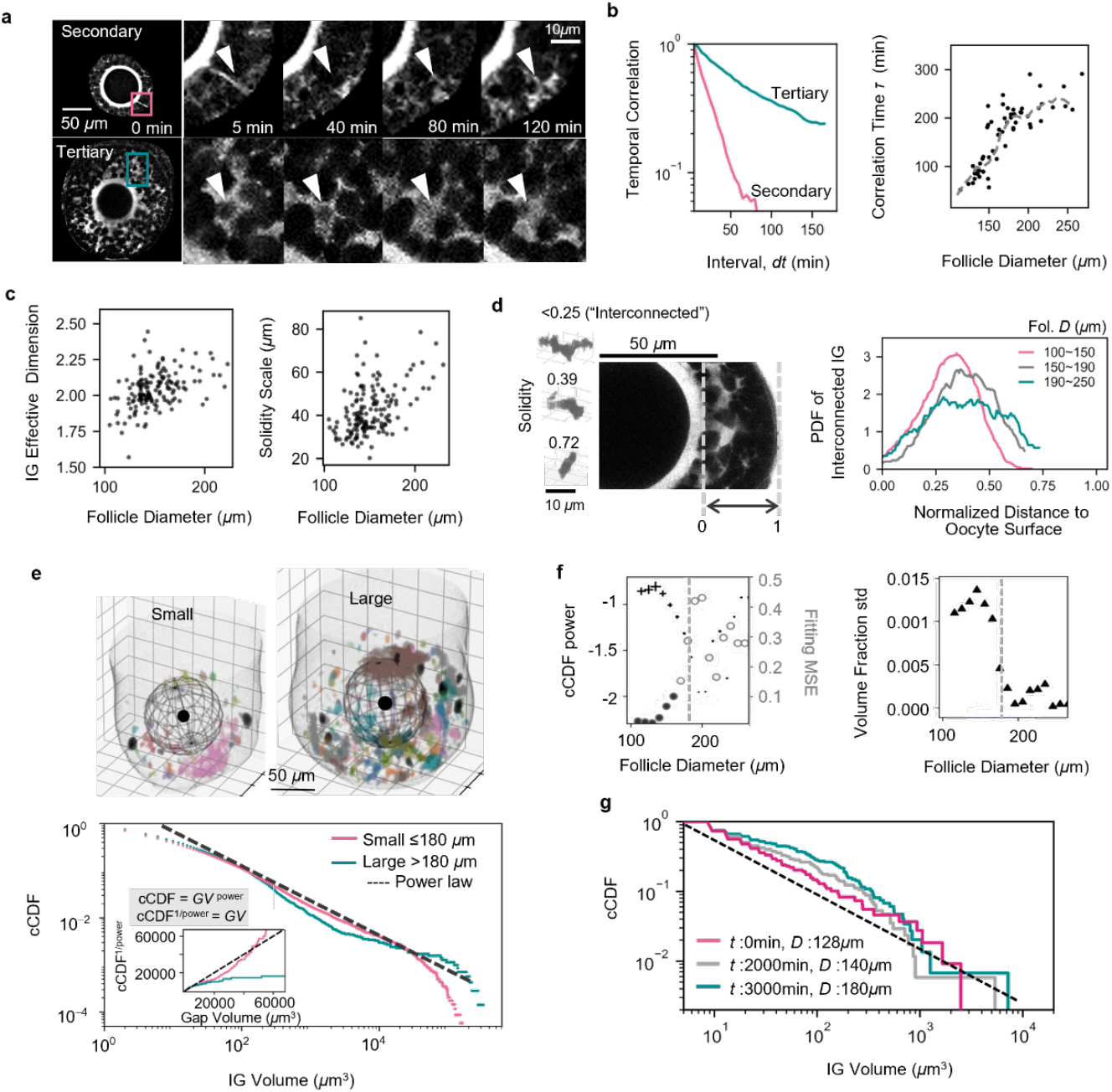
Regime transition underlies fluid growth during secondary-tertiary follicle development. **a**, Representative images of secondary (top) and tertiary (bottom) follicles stained with FITC and tracked over 5 h, showing zoomed-in regions containing IGs at different time points. See Supplementary Video 3 and 4. **b**, Left: Representative plot of temporal correlations of IG signals in secondary (pink) and tertiary (green) follicles. Right: Plot of correlation time of IG signals of each follicle against follicle diameter (N =14 , n = 55). **c**, Plots of IG effective dimension (left, Pearson *R* = 0.432, *p* = 2×10^-8^) and solidity scale (right, Pearson *R* = 0.472, *p* = 8×10^-10^) derived from each follicle against follicle diameter. **d**, Left: Image of interconnected (solidity <0.25) IGs near the oocyte (left); Right: plot of average probability density function (PDF) of interconnected IGs over the normalised distance from oocyte surface, for follicles of different size range. **e**, Top: Representative IG morphology in small (left) and large (right) follicles. Each colour in the 3D-rendered image represents a distinct IG. Bottom: Pooled plot of complementary cumulative distribution function (cCDF) of IG volume in follicles smaller (pink) and larger (green) than 180 µm. Inset: Corresponding plot of reciprocal power of cCDF against IG volume. **f**, Left: Plot of fitted cCDF power (left axis) and its corresponding mean square error (MSE, right axis) against follicle diameter. Right: Plot of standard deviation of IG volume fraction sagainst follicle size. Dotted lines represent 180 µm. **g**, Plot of IG cCDF against volume in the follicle shown in Fig. 1e, showing their temporal evolution with increasing follicle size *D*. N =15, n=153 follicles, 41764 gaps for **c**-**f**.

Next, we examined the morphology of IGs across follicle size. To quantify the detailed morphological features of IGs, we measured the increase in IG volume *V* with its length *L* in each follicle, and extracted the scaling factor as an effective dimension (see Methods, Extended Data Fig. 3a). An effective dimension close to 3 means that the IG expands in a spherical manner in three dimensions (3D), while a value close to 1 means that the IG expands effectively in a one-dimensional manner, like a thread. With this, we observed an increase in effective dimension with follicle size (Fig. 2c, left), suggesting that the IGs are transitioning from a network-like topology to more spherical-like cavities during follicle development. Similar trend was observed when we quantified IG solidity (see Methods), which reflects how convex (close to 1) or fractal-like (close to 0) a shape is^15^. By defining a solidity scale as a typical length scale beyond which the IG solidity saturates at a low value (Extended Data Fig. 3b), we found that this solidity scale correlates with follicle size (Fig. 2c, right), suggesting that the IGs are more fractal-like in secondary follicles. Additionally, we found that large interconnected IGs are located close to the oocyte in small follicles but were situated further away from the oocyte with increased follicle size (Fig. 2d, Extended Data Fig. 3c). This was not observed for the more ‘solid’ IGs (Extended Data Fig. 3d), suggesting that only the localisation of interconnected IGs depends on the stage of follicle development.

The transition from interconnected to more compact IGs during secondary-tertiary follicle development implies a global change in the lumen configurations. To investigate this, we next analysed the IG size distribution to extract global features of fluid dynamics. In plotting the pooled complementary cumulative distribution function (cCDF) of IG volume (Fig. 2e), we observed a near power-law distribution of IG volume in secondary follicles which span across more than two orders of magnitude. This indicates that the IGs in the small follicles do not have a characteristic length scale. By contrast, we observed a bimodal profile in tertiary follicles in which small IGs with sizes less than 10 μm coexist with very large IGs (∼50 μm). This transition in IG statistics is not exclusive to isolated follicles, as we also observed a similar trend in ovarian microtissues (Extended Data Fig. 4a, b). The transition in IG characteristics is most pronounced at 180 μm, where we observed a striking increase in the mean square error (MSE) of power-law fitting (Fig. 2f, left) and a sharp decrease in the standard deviation of IG volumes (Fig. 2f, right), further supporting the notion that the IGs in secondary follicles exhibit high volume fluctuations. To confirm our finding, we directly tracked IG distribution in single follicles over time. We found that while the initial IGs in a secondary follicle followed a power-law distribution, with time, the IG distribution is characterised by a broad exponential profile for small gaps and a few large gaps at the tail (Fig. 2g), consistent with our population-based study.

In summary, our results suggest the existence of two distinct regimes of IGs during development from the secondary to tertiary follicle stage. In secondary follicles, IGs are highly dynamic and interconnected, exhibiting large fluctuations in volume and shape, while in tertiary follicles, IGs are more stable and morphologically compact, with a few large IGs dominating lumen growth, reminiscent of the transition from near-critical behaviour to phase separation^16^.

### GCs are highly dynamic and show increased cell death during follicle development

We next explored how changes in GC activities may contribute to the formation and organisation of IGs. To this end, we first tracked GC dynamics by quantifying the mean squared displacement (MSD) of GCs in H2B-mCherry follicles (Fig. 3a, b and Supplementary Video 5). MSD for GCs increases exponentially with lag time, exhibiting diffusive behaviour at a long timescale (>1 h) (Fig. 3b), indicating that the GCs are fluid-like at the timescale of follicle development. This fluid nature of GC dynamics is robust during development, as shown by a similar value of MSD power across a wide range of follicle size (Extended Data Fig. 5a). For cell motion on short timescale (<1 h), where MSD power is much lower, GCs are caged by the surroundings neighbours. Using label-free quantitative phase imaging (QPI) technique^17^ and particle image velocimetry (PIV) (Extended Data Fig. 5b and Supplementary Video 6), we found a median instantaneous velocity of 0.063 µm/min for the GCs (Extended Data Fig. 5c, d). This indicates that GCs exhibit intrinsic random motions at short timescales, which may allow the cells to achieve fluid-like rearrangements at longer timescales.

**Figure 3:**
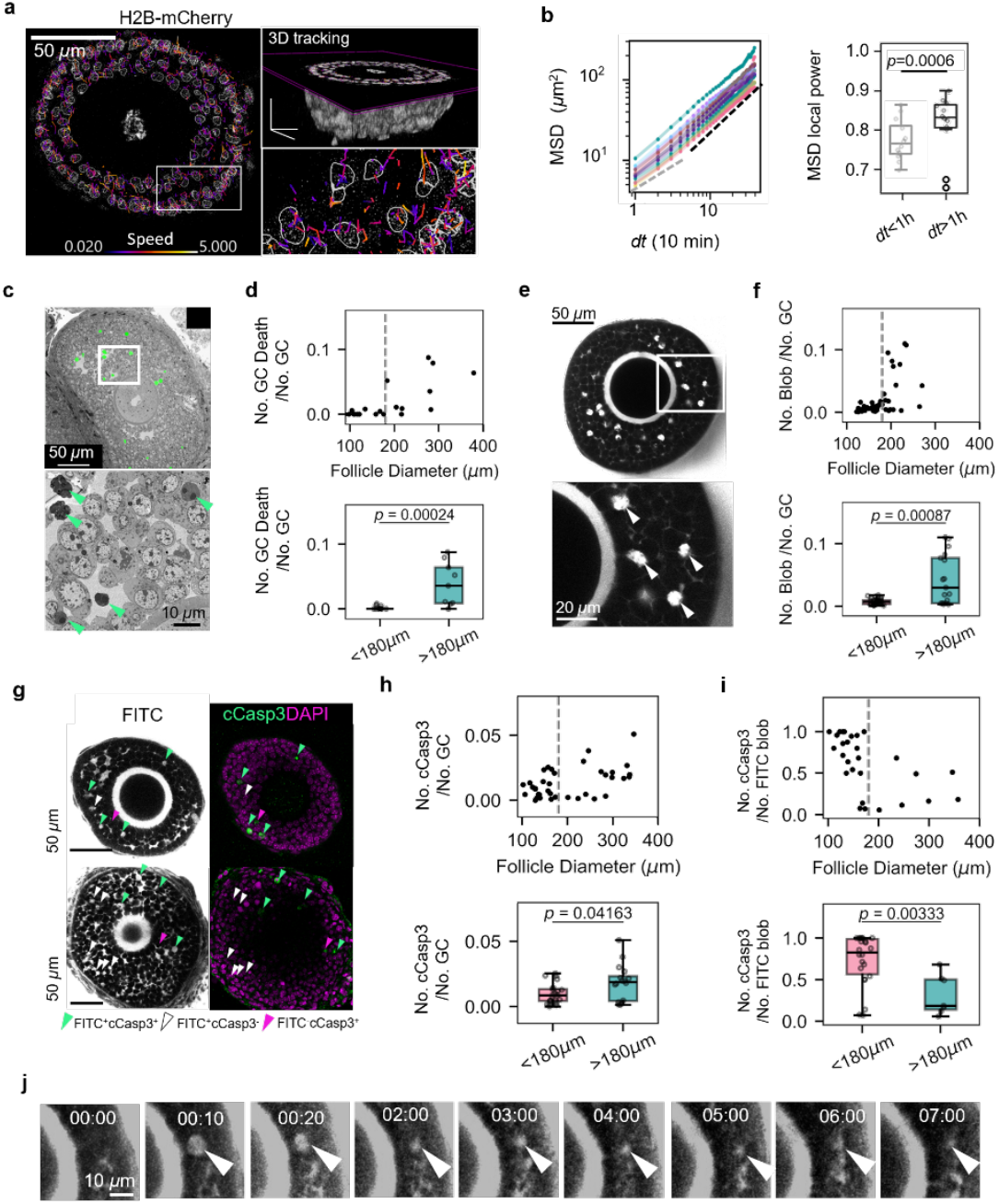
GCs are dynamic and show increased cell death in tertiary follicles. **a**. Representative image of a follicle expressing H2B-mCherry under timelapse imaging over 16 h. Colour coded tracks show the speed of each segmented nucleus within the follicle over time. See Supplementary Video 5. **b**, Left: Plots of mean-squared displacement (MSD) against time interval. Right: Boxplot of MSD local power for time intervals of less and more than 1 h (N = 2, n = 12). **c**, Representative images of a follicle imaged with SEM. Inset: Zoomed-in image of white box, showing dead GCs (green arrowheads). **d**, Top: Plot of normalised number of dead GCs against follicle diameter. Bottom: Boxplot of normalised number of dead GCs for follicles smaller and larger than 180 µm (n = 22 follicles). **e**, Representative image of FITC blobs (white arrowheads) in isolated follicles stained with FITC. Inset: Zoomed-in image of white box. **f**, Top: Plot of normalised number of FITC blobs against follicle diameter. Bottom: Boxplot of number of normalised FITC blobs for follicles smaller and larger than 180 µm (N = 11, n = 47). **g**, Representative image of follicles smaller and larger than 180 µm, showing FITC, nuclei (grey), cCASP3 (green) and actin (magenta). Green arrowheads show FITC blob co-localised with cCASP3 while white arrowheads show FITC blob without cCASP3 co-localisation. **h**, Top: Plot of normalised number of cCASP3 stained GCs against follicle diameter. Bottom: Boxplot of normalised number of cCASP3 stained GCs for follicles smaller and larger than 180 µm (N = 6, n= 37). **i**, Top: Plot of ratio of FITC blobs stained with cCASP3 against follicle diameter. Bottom: Boxplot of ratio of FITC blobs stained with cCASP3 for follicles smaller and larger than 180 µm (N = 5, n = 29). **j**, Representative images of a FITC blob (white arrowhead) tracked over time, showing disappearance of a blob and merging with the IG. See Supplementary Video 8.

Since cell death is known to contribute to lumen formation^18^, we quantified the extent of cell death during follicle development. Using electron microscopy and fluorescence imaging of FITC-stained follicles, we showed that the number of cell death events was close to zero in secondary follicles, but started to increase in the tertiary follicles (Fig. 3c-f). We confirmed the FITC-positive blobs were dead cells, by showing that it co-localised with cleaved caspase 3 (cCasp3), an apoptosis marker, which can be induced in staurosporine treated follicles (Fig. 3g, Extended Data Fig. 6a). When stained with cCasp3, we observed fewer apoptotic events in secondary follicles but a gradual increase in follicles larger than 180 µm (Fig. 3h), consistent with our findings from electron microscopy and FITC staining studies. Since the percentage of apoptotic events were lower than the total cell death quantified, we investigated the proportion of blobs that were apoptotic. We observed that in small follicles, most blobs co-localised with cCasp3 signals (FITC^+^cCasp3^+^), while in tertiary follicles, there was an increased number of blobs without cCasp3 signals (FITC^+^cCasp3^-^) (Fig. 3g, i). To note, some cCasp3 signals were not co-localised to any blobs (FITC^-^cCasp3^+^), suggesting early apoptotic events. Altogether, these data suggest that the early initiation of GC death may be triggered by apoptosis, but other cell death pathways may be implicated as well.

To better map out the temporal dynamics of cell deaths without incurring phototoxicity, we performed timelapse imaging on isolated follicles with QPI. In follicles of all sizes, we observed apoptotic events characterised by high refractive index (RI), possibly due to chromatin condensation during apoptosis. In tertiary follicles, we occasionally observed low RI structures that are highly spherical and have a size similar to that of GCs. They typically appeared for about 15-30 min before vanishing into the background (Extended Data Fig. 6b and Supplementary Video 7). This structure is reminiscent of lytic cell death events regulated by necroptosis^19^. Indeed, FITC-labelled blobs underwent swelling and rupture (Extended Data Fig. 6c), similar to the described characteristics of necroptotic cell death^20^. At longer times, these blobs lost their integrity and were “sheared” into larger IGs (Fig. 3j, Supplementary Video 8). This data suggests that GC death may contribute to overall IG volume increase at the onset of secondary-tertiary follicle transition.

Apart from GC death, cell polarity can also drive *de novo* lumen formation^10,21^. To this end, we examined markers related to apical polarity proteins. Remarkably, we found that the GCs organise into rosette structures with ECM-filled cavity in the centres (Extended Data Fig. 7a-f, k). These rosette GCs exhibit basal markers like integrin (Extended Data Fig. 7c) at the cavity-facing side of the cells, and apical markers such as Golgi apparatus, tight junction (ZO-1), endosomal vesicle and microvilli (acetylated-alpha tubulin) at the opposite end (Extended Data Fig. 7d-g), suggesting that the GC rosettes are basolaterally polarised and not likely generated by apical polarity-driven mechanisms, as observed in epithelial cells^10,21^. During follicle development, we noticed that the size of the rosette ECM centres became smaller and more speckled with follicle size (Extended Data Fig. 7h), and their number appeared to peak when the follicles grow to 180 µm (Extended Data Fig. 7i). Spatially, the rosettes appeared closer to the oocyte in smaller follicles but became more evenly distributed in larger follicles (Extended Data Fig. 7j). Of note, we observed a co-localisation of FITC signals with CNA35, which labels pan-collagen in the ECM-enriched rosette centres. This suggests that GC rosettes do contribute to small IGs in follicles (Extended Data Fig. 7k).

### IG transition is accompanied by spatiotemporal changes of GC junction mechanics and motility

We next investigate changes to junctional dynamics between GCs during secondary-tertiary follicle development. Focusing on the basal GCs sitting at the outermost layer of the follicle, we observed frequent occurrence of cell neighbour exchanges (T1 transition events), which has been reported to reflect tissue fluidity^22^. We found that the T1 transition events in the basal GCs increases with follicle size, indicating increased tissue fluidisation during follicle development (Fig. 4a, b and Supplementary Video 9). To understand this phenomenon at cellular level, we stained the GCs with N-cadherin (N-cad), the primary cell-cell adhesion molecules connecting the GCs^23^ (Fig. 4c and Extended Data Fig. 8a). We found that the N-cad continuity decreases with follicle size (Fig. 4c), which validates our use of N-cad continuity as a proxy for cell-cell junctional strength in GCs. In addition, the radially projected junctional length of basal GCs decreases with follicle size (Fig. 4d and Extended Data Fig. 8b). Overall, these data indicate an overall reduction in the basal GC intercellular tension during follicle growth, consistent with our observation of increased T1 transition in basal GCs during this period.

**Figure 4:**
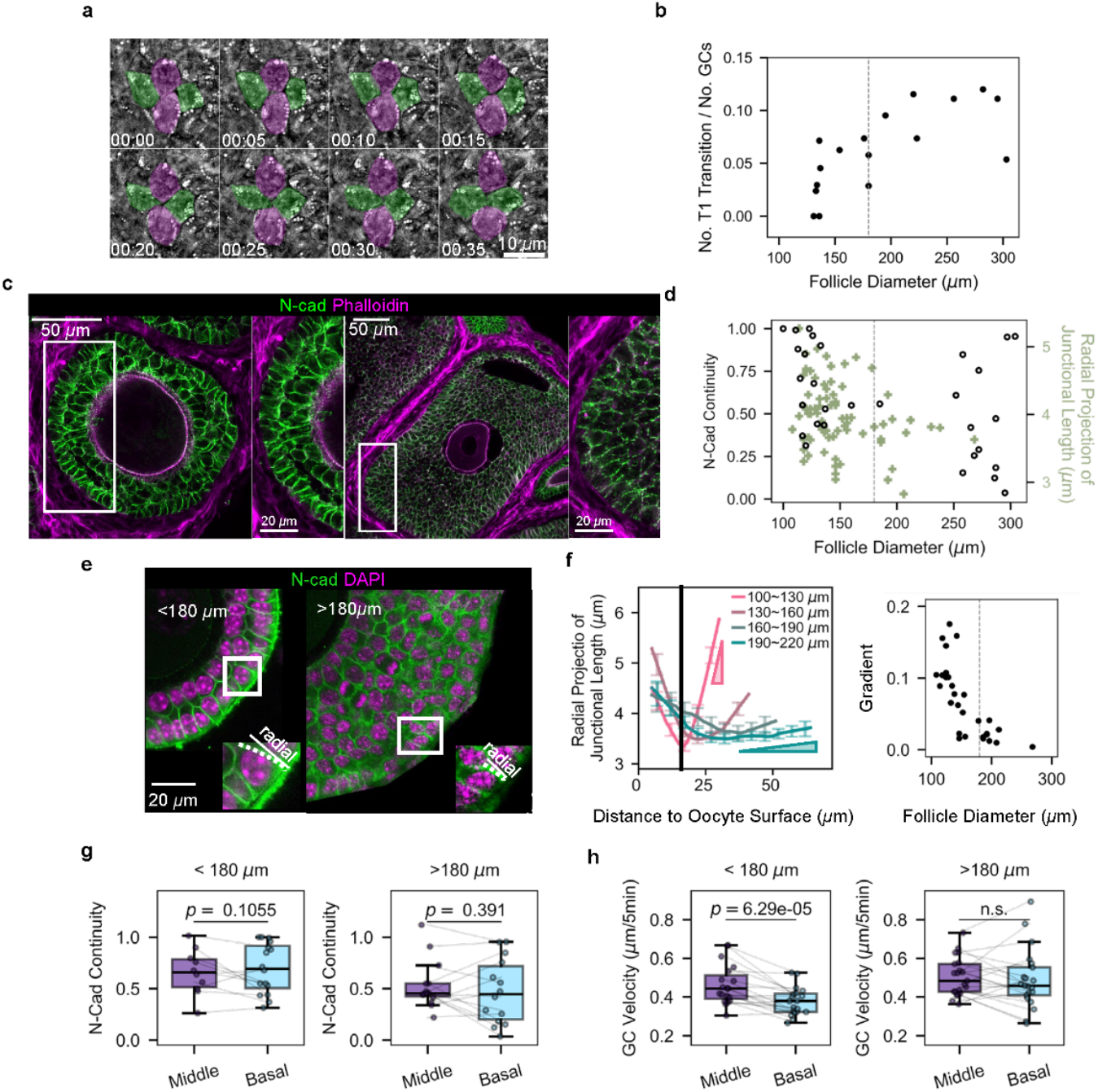
GCs exhibit spatial gradients in cell-cell adhesion and cellular dynamics. **a**, Representative images of GC rearrangements (termed T1) with time, captured by quantitative phase imaging. Green and magenta colours indicate representative T1 transition. See Supplementary Video 9. **b**, Plot of normalised T1 events for basal GCs against follicle size. Pearson correlation *R =* 0.072, p-value=0.001 (N = 6, n = 17). **c**, Representative images secondary (left) and tertiary (right) follicle in tissue slice stained with N-cad (green) and Phalloidin (magenta). Zoomed in image of white box shows difference in N-cad continuity. **d**, Plots of N-cad continuity along basal GC lateral junctions, against follicle size (black circle, Pearson correlation *R* = -0.4, p-value = 0.014) (N = 2 , n = 30), and radial projections of basal GC lateral junction length against follicle size (green cross, Pearson correlation *R* = -0.280, p-value=0.008) (N = 3, n = 29). **e**, Representative images of isolated follicles stained with DAPI (magenta), N-cad (green), for follicles smaller and larger than 180 µm. Right: Zoomed in image of white box showing projection of each cell-cell lateral junction onto the radial direction (dotted line). **f**, Left: Plots of radial projections of GC junctional length against their distance to the oocyte surface, pooled from different follicle size. Right: Plots of linearised gradients of projection lengths between the middle and basal GC layers, against follicles size (N = 3, n = 29). **g**, Boxplots of N-cad continuity for the middle and basal GC layers, for follicles smaller (left) and larger than 180 µm (right) (N = 2, n = 30). **h**, Boxplots of GC median velocity for the middle and basal GC layers, for follicles smaller (left) and larger than 180 µm (right) (N = 13, n = 40). All dashed lines indicate 180 µm.

Noting that the GC shape varies spatially across the follicle (Fig. 4c, e), we quantified the radial projection of each GC junctional length across the entire follicle (see Methods) and extracted its gradient from the middle to basal GC layers (Fig. 4f). For secondary follicles, we observed a steep gradient in the radially projected junctional length, which gradually diminished with increased follicle size (Fig. 4e, f). Furthermore, in tertiary follicles, the ratio of radial to tangential projected length of GC junctions approaches 1 in both the middle and basal GCs, suggesting that the GCs become uniformly isotropic in shape across the entire follicle (Extended Data Fig. 8c). Focusing on N-cad expression, we found no difference in N-cad junctional continuity between the middle and basal GC layers in both secondary and tertiary follicles (Fig. 4g). Taken together, our findings reveal the presence of a gradient of intercellular tension between the middle and basal GC layers, particularly in the secondary follicles, which become diminished as the follicles grow to become tertiary follicles.

Given that the GCs also display instantaneous motions (Fig. 3a, b), we next examined if there exist any spatial variations in GC motions within the follicle. Using QPI and PIV, we found that the middle GC velocity was significantly higher than that of the basal GCs in the secondary follicles (Fig. 4h). However, this difference disappeared in the tertiary follicles, as the basal GCs became significantly more dynamic (Fig. 4h, right and Extended Data Fig. 9d). This trend is similar to the observed decrease in the gradients of radially projected length of basal GCs across all follicle size (Fig. 4f).

### Binary fluid model recapitulates IG regime transition during follicle development

To get systematic insights on the mechanisms underlying IG dynamics, we developed a physical model that integrates our experimental findings presented thus far. Given that GCs are fluid-like on the timescale of follicle development, we modelled the system as a binary fluid mixture (GCs and IGs) in a closed space. We hypothesized that IG dynamics arises from interactions between IGs and GCs, and implemented the system as an Ising model with conserved elements^24^ (see Supplementary Information). Briefly, the dynamics is governed by local exchanges between nearest neighbours to minimize cell-cell coupling (*J*), which competes with an effective temperature (*T*) representing the noise of exchange. A large *J* reflects strong interfacial tension that prevents cell-lumen mixing, resulting in phase separation. Conversely, a high *T* allows cells to overcome cell-cell adhesion, promoting mixing between cells and lumen. The ratio of *J* over *T* is therefore a key control parameter of the system. From experiments, we attribute *J* to the combined effect of GC junctional length and N-cad continuity (Fig. 4c, d), while *T* is represented by the instantaneous velocity of GCs (Fig. 4h). Our data (Fig. 4f-h) implies a presence of a spatial gradient in *J*/*T*, with the lowest values near the oocyte and the highest at the basal GCs, which was incorporated into our simulation (Fig. 5a, left).

**Figure 5:**
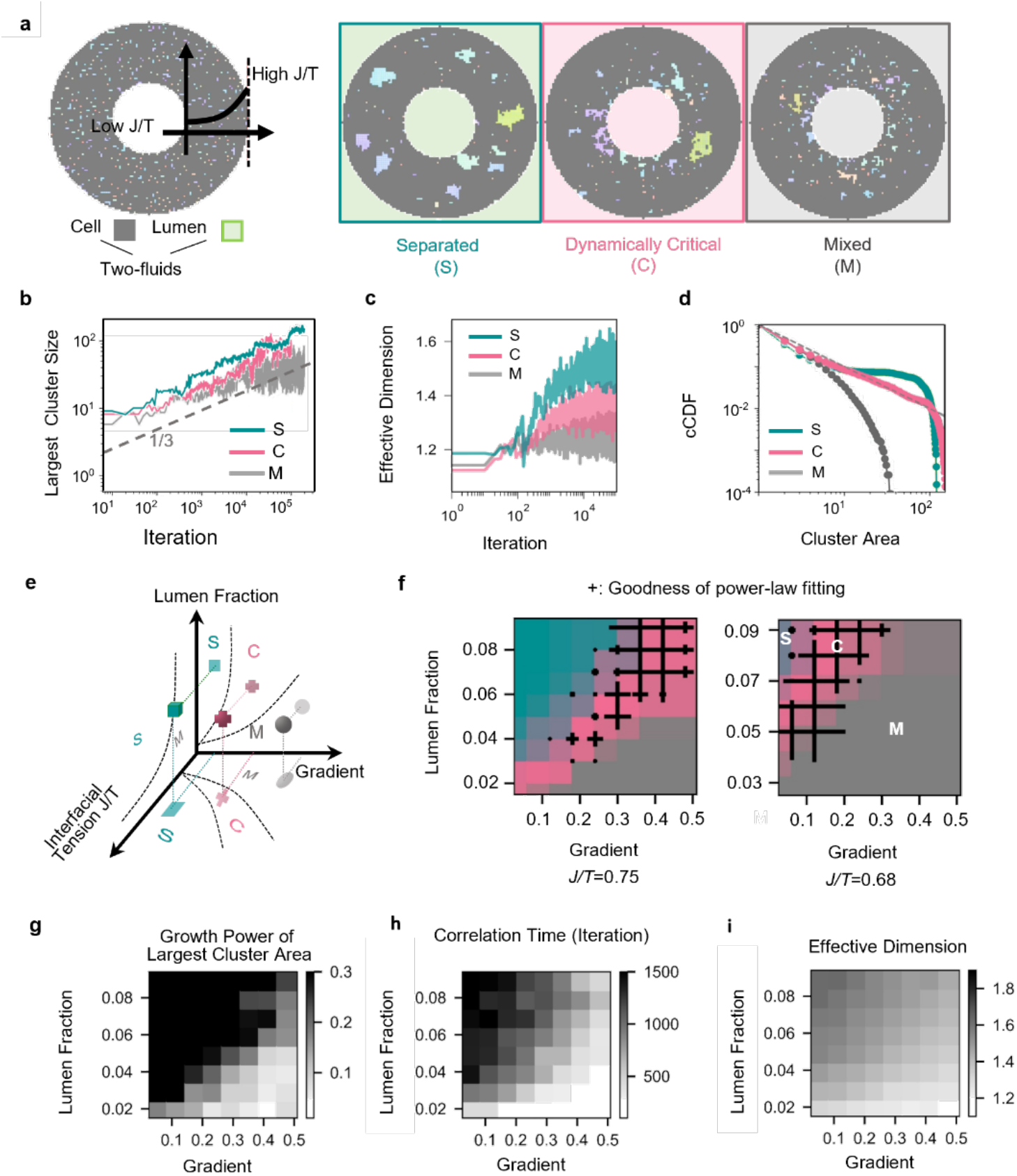
Binary fluid model with interfacial tension gradient recapitulates IG transition during follicle development. **a**, Illustration of a 2D lattice setting with fixed boundaries representing oocyte surface and basal surface. Each lattice is assigned as 1 for cell or -1 for lumen randomly at initial step and the effective interfacial tension *J*/*T* has a gradient from inner to outer GC layer; Representative snapshots for three typical steady states, separated (S), dynamically critical (C) and mixed (M). **b**, Illustration of 3D phase diagram with three control parameters: lumen fraction, interfacial tension *J*/*T* and the gradient of *J*/*T*. **c**, Plot of representative growth curves for largest gap size with time. **d**, Plot of representative growth curves for effective gap dimension with time. **e**, Representative cCDF profiles of gap area for the three distinct phases. **f**, 2D phase diagram with a global *J*/*T* fixed at 0.75 (left) and 0.68 (right). The size of cross corresponds to the goodness of fit to the power law for each set of control parameters. **g-i**, Maps corresponding to a global *J*/*T*=0.75. Colours indicate the growth power of largest gap area (**g**), temporal correlation scale in the unit of iteration step (**h**), and effective gap dimension (**i**), respectively.

After searching through the parameter space, we observed three steady states being reached, namely the separated (S), dynamically critical (C), and mixed (M) states. Their representative lumen configurations in 2D are presented (Fig. 5a, right and Supplementary Video 10 and 11). Separated state is characterized by the nucleation of stable lumen clusters uniformly scattered across the space, while the mixed state contains many small isolated and transient lumens. In dynamically critical state, we observed large lumen clusters near the oocyte, and small gaps with fuzzy boundaries. We investigated the dynamic features of the three states and found that the size of largest clusters in the S and C states grew with time (over 10000 iterations) to the power of 1/3, while the largest clusters in the mixed state did not increase (Fig. 5b). We further quantified the effective dimension of cluster, using the same methods as for experiments (Fig. 2c, left). While the effective dimension grew and approached the space dimension for the case of S state, they only reach a saturation value below the space dimension in the C and M states (Fig. 5c). Notably, as the system approached steady state, the cluster size distribution revealed three distinct profiles: S state characterised by a bimodal profile with a peak for largest clusters, M state showing a fast exponential decay, and the C state exhibiting near power-law distribution with a cutoff at system size (Fig. 5d). The lumen configuration and cCDF profiles for C state and S state qualitatively recapitulate the observed near-critical regime in secondary follicles and the stable growth regime in tertiary follicles (Fig. 2d-e).

As simulation on 2D and 3D lattice show qualitatively the same results (see Supplementary Information for 3D simulation), we explored the full phase diagram with 2D lattice for the sake of computational efficiency. The parameters governing the IG steady states are *J*/*T*, gradient of *J*/*T*, and lumen fraction. Together, they allow us to construct a 3D phase diagram (Fig. 5e). The phase boundary between dynamically critical state and the other two states are determined by a sharp enhancement in the goodness of power-law fitting of cCDF (Fig. 5f). Of note, the dynamically critical state only exists for non-zero gradient of *J*/*T*. From the 2D projection of the phase diagram, we found that the dynamically critical state exists between the mixed and separated state, and its region expands with the gradient of *J*/*T*. Reducing global *J*/*T* effectively expands the region of M state, as expected when the fluids are highly dynamic or interfacial tension is low. From our phase diagram, we infer that the transition from secondary follicle (C state) to tertiary follicle (S state) can be triggered by a weakening of *J*/*T* gradient, an increase in lumen fraction, or a strengthening of a global *J*/*T*. Experimentally, our data are consistent with the former two predictions (Fig. 1c-d, 4f). However, our data (Fig. 4g, h) appear to imply a decrease in global *J*/*T*. We propose that this primes the system to slightly deviate from the critical regime, such that only a minimal number of rapidly growing dominant lumens are observed, as found in tertiary follicles.

We next generated output from simulation in terms of the largest cluster growth power (Fig. 5g), correlation time of lumen (Fig. 5h) and effective dimension (Fig. 5i). As shown from the maps, a transition from C to S corresponds to an increase in correlation time and effective dimension, in line with experimental data (Fig. 2b-d). However, the growth power of the largest cluster does not change during the transition from C to S phase, as both follow roughly a power of 1/3. The growth of the largest cluster size in the dynamically critical state exhibits more temporal fluctuations than the separated state, which matches the observed dynamics (Fig. 1g). We next incorporated additional non-conservative elements, such as cell proliferation and death, into our model (see Supplementary Information). By integrating information on the spatial distribution of GC proliferation and death in secondary and tertiary follicles (Extended Data Fig. 9a-d), our model (Extended Data Fig. 9e) predicts that the three steady states (S, C, and M) remain unaffected, as long as the growth of lumen creation is sufficiently slow (see Supplementary Information for further discussion), which matches our experimental data (Fig. 1g, purple crosses). In summary, our minimal binary fluid model reveals that lumen growth is primarily driven by the spatiotemporal change of cell-lumen interactions (*J/T*), with cell death and proliferation playing a more secondary role in dictating the lumen critical behaviour.

### Perturbing *J/T* and fluid fraction impact lumen critical behaviour and follicle growth

In our model, the distinct phases of lumen in follicles are governed by the overall *J/T* and gradient of *J/T* across GC layer. To test the model, we perturbed the interfacial tension *J* by disrupting granulosa cell-cell adhesion, using ADH-1, an N-cad antagonist. To validate its effect, we first measured changes in the contact angle and cell-cell distance between two GCs in a doublet, which had been shown to reflect cell-cell adhesion strength^25,26^. Indeed, ADH1-treated doublets showed decreased contact angles and increased cell-cell distance (Fig. 6a), reflecting reduced cell-cell adhesion following ADH-1 treatment. The effect was also demonstrated in follicles, where ADH-1 treatment led to reduced N-cad continuity of basal GCs, decreased gradient of radial junctional length, and increased T1 transition events in basal GCs (Fig. 6b-d and Supplementary Video 12). By contrast, ADH-1 did not result in any change in the middle GC velocity (Fig. 6e and Supplementary Video 13), suggesting negligible impact on *T*. Together, these data indicate that ADH-1 treatment led to reduced *J/T*. We next examined the impact of ADH-1 on IG phase. We observed a decrease in temporal correlation (Fig. 6f) and effective dimension (Fig. 6g) following ADH-1 treatment, suggesting IG fragmentation due to higher fluid dynamics. Furthermore, direct tracking of IGs in follicles treated with ADH-1 over 72 h showed that the IGs transitioned from a power-law to an exponential distribution during this period (Fig. 6h). Overall, our data indicate that the IGs are becoming more dynamic, in line with our model prediction that decreased *J/T* favours the lumen to exist in the mixed phase (Fig. 5e).

**Figure 6:**
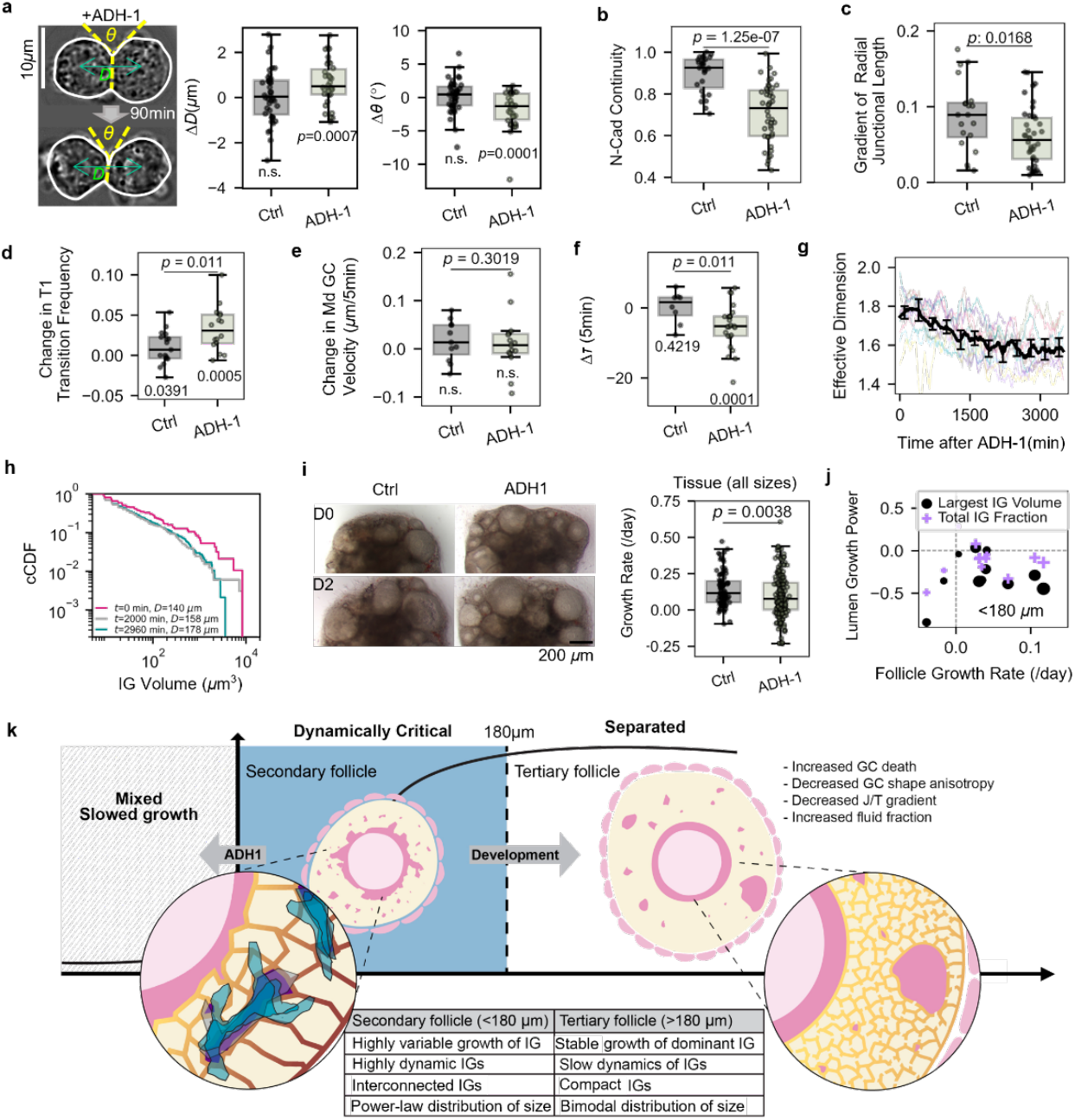
Perturbing *J*/*T* impacts lumen criticality and follicle growth. **a**, Left: Representative images of GC doublet at 0 min (top) and 90 min (bottom) after ADH-1 treatment, showing cell-cell distance (*D*) and contact angle (*θ*). Right: Boxplots of changes in *D* (middle) and contact angle (right) for control (N = 2, n = 40) and ADH-1 after 90 min of treatment (N = 3, n = 34). **b**, Boxplot of N-cad continuity for isolated follicles in control (N = 2, n = 44) and ADH-1 conditions (N = 2, n = 77). **c**, Boxplot of radial projection of basal GC lateral junction length for isolated follicles in controls (N = 2, n = 29) and ADH-1 conditions (N = 2, n = 46) **d**, Boxplot of normalised number of T1 transition events in control-DMSO (N = 2 n = 17) and ADH-1 conditions (N = 3, n = 16), **e**, Boxplot of change in middle GC velocity for follicles in control-DMSO (N = 2, n = 10) and ADH-1 conditions (N = 3, n = 15). **f**, Plot of change in the temporal correlation timescales of IG against follicle diameter in isolated follicles in control (N = 2, n = 8) and post ADH-1 treatment (N = 6, n = 25). **g**, Plot of IG effective dimension against time after treatment of ADH-1 (N = 2, n =12). **h**, Plot of temporal evolution of cCDF for a representative follicle after ADH-1 treatment. **i**, Boxplots of follicle growth rates for microtissues in controls and ADH-1, tracked over 5-7 days (right), with representative images shown (left) (N = 2, n = 49 (control), 78 (ADH-1)). **j**, Plot of growth power of the largest IG volume (black circle) and total IG fraction (purple cross) against mean follicle growth rate for isolated follicles tracked over 3-4 days (N = 2, n = 12). **k**, Summary schematic showing how changes in IG characteristics, cell death and GC junctional mechanics contribute to lumen criticality transition during secondary-tertiary follicle development.

To investigate the functional impact on follicles when the IGs are challenged to remain in the mixed phase, as in the case of ADH-1 treatment, we quantified the overall follicle and IG growth kinetics. We found that while ADH-1 treated follicles continued to grow, they experienced a slower growth rate, as observed in microtissues (Fig. 6i). Notably, ADH-1 treated follicles exhibit zero or negative growth in the IGs (Fig. 6j), suggesting that perturbing the critical phase of IGs can negatively impact the overall follicle development. To investigate how GC apoptosis impacts lumen fraction and follicle growth, we treated follicles with Z-VAD-FMK, a broad-spectrum inhibitor of caspases. In this case, the cCDF distribution for the treated follicles assumed a more exponential profile rather than power-law behaviour (Extended Data Fig. 9f), indicating a propensity for the system to move away from the S state (Fig. 5f). Notably, Z-VAD-FMK-treatment led to decreased growth rate for tertiary follicles (Extended Data Fig. 9g), further supporting our conclusion that perturbing lumen growth can impact functional growth of follicles.

## DISCUSSION

Recent studies across multiple organisms have highlighted the importance of lumen in regulating developmental programs^27^. For instance, the change in fluid distribution and cellular organisation is essential for morphogen distribution in zebrafish embryo development^25^. Lumen can also act as a signalling hub to drive cell differentiation and cell shape changes around the lumen^28,29^. In this study, we report, for the first time, that the *de novo* lumen formation during murine folliculogenesis is characterised by the emergence of highly dynamic and interconnected fluid pockets close to the oocyte, followed by the gradual coalescence into larger, stably growing gaps across the entire follicle (Fig. 6k). This behaviour, which resembles classical phase separation, appears to be regulated by increased GC death and changes in granulosa cell-cell adhesions. This supports the notion that the release of cytoplasmic materials and DNA may act as a source of osmotic potential to regulate lumen fraction, as speculated in past studies^9^. We found that in secondary follicles, the presence of a gradient of interfacial tension regulated by N-cad adhesions between GCs, helps to drive the fluid globally towards the oocyte, as predicted by our minimal binary fluid model. Our findings echo other studies showing that intercellular tension guides lumen morphology and position at tissue scales^25,30,31^. Importantly, this *gradient* of intercellular tension is crucial in maintaining the IGs in a *sustained* critical state over an extended period of time (days), in contrast to classical phase transitions in non-living systems (e.g. liquid-gas transition), where the critical point is highly sensitive to external perturbations and is therefore not observed easily.

Given that the oocytes are known to grow in size till ∼180 µm^32^, we propose that the highly dynamic IGs near the oocyte during this phase may facilitate rapid exchange of nutrients in the oocyte microenvironment, thereby supporting oocyte maturation. Future work to track fluid dynamics within the follicle using tracer particles, and functional studies on oocyte transcriptome following fluid perturbations, may shed light on this aspect. Of note, the observed transition in IG phase near 180 µm coincides with an important biological transition, where the follicles are becoming FSH-dependent^33^. This may trigger the observed increase in GC death and proliferation, which can contribute to increased lumen fraction, GC rearrangements, and effective phase separation.

Interestingly, we observed that deformed follicles in microtissues typically show faster lumen coalescence and exit of near-critical state (Extended Data Fig. 10a, b), suggesting that extrinsic mechanical force may influence gradient of interfacial tension and luminogenesis. Indeed, growing follicles at the ovarian cortex typically experience greater deformation, which may prime these follicles to undergo earlier lumen expansion and follicle development. This constitutes an exciting perspective where follicle position, mechanics and morphogenesis are tightly coordinated to give rise to the robust selective growth of follicles and eventual ovulation.

## Acknowledgements

We thank Ling Wang (Prevedel group, EMBL Heidelberg) for technical assistance; Dang Hairuo (Ellenberg group, EMBL Heidelberg) for providing ovarian tissues from 48 weeks R26-H2B-mCherry mice (Identifier: CDB0239K). We thank Jacques Prost, Daniel Riveline and Raymond Rodgers for discussion and feedback on the manuscript. The Chan lab is supported by the Ministry of Education under the Research Centres of Excellence programme through the Mechanobiology Institute and the Department of Biological Sciences at the National University of Singapore, the Ministry of Education Tier2 grant (T2EP30222-0026), National Research Foundation Mid-size Grant (NRF-MSG-2023-0001) and the Bia-Echo Asia Centre for Reproductive Longevity and Equality (ACRLE) at the National University of Singapore. C.J.C. acknowledges the support of the Singaporean Teaching and Academic Research Talent Inauguration Grant (START). C.B. and R.P. acknowledge funding from the EMBL and an ERC Consolidator Grant (grant no. 864027, Brillouin4Life). T.H. acknowledges the partial support by the National Science and Technology Council, Taiwan (NSTC113-2112-M-001-051-MY3).

## Author contributions

Y.L., K.W.L., T.H., C.J.C. conceptualized the project. K.W.L., Y.L. and C.J.C. designed experiments. K.W.L., A.B., J.Y.K.T., X.P.J.T., X.L., B.H.N., T.B.L. and C. B. performed the experiments. K.W.L., Y.L., J.Y.K.T., X.P.J.T. and X.L. analysed the data. K.W.L., Y.L. and C.J.C. wrote the manuscript. T.H. helped with the interpretation of the results and commented on the manuscript. R.P. and I.B. provided support in Brillouin microscopy and SEM, respectively. C.J.C acquired funding and supervised the study.

## Competing interests

The authors declare no competing interests.

## EXTENDED DATA FIGURES

**Extended Data Fig 1:**
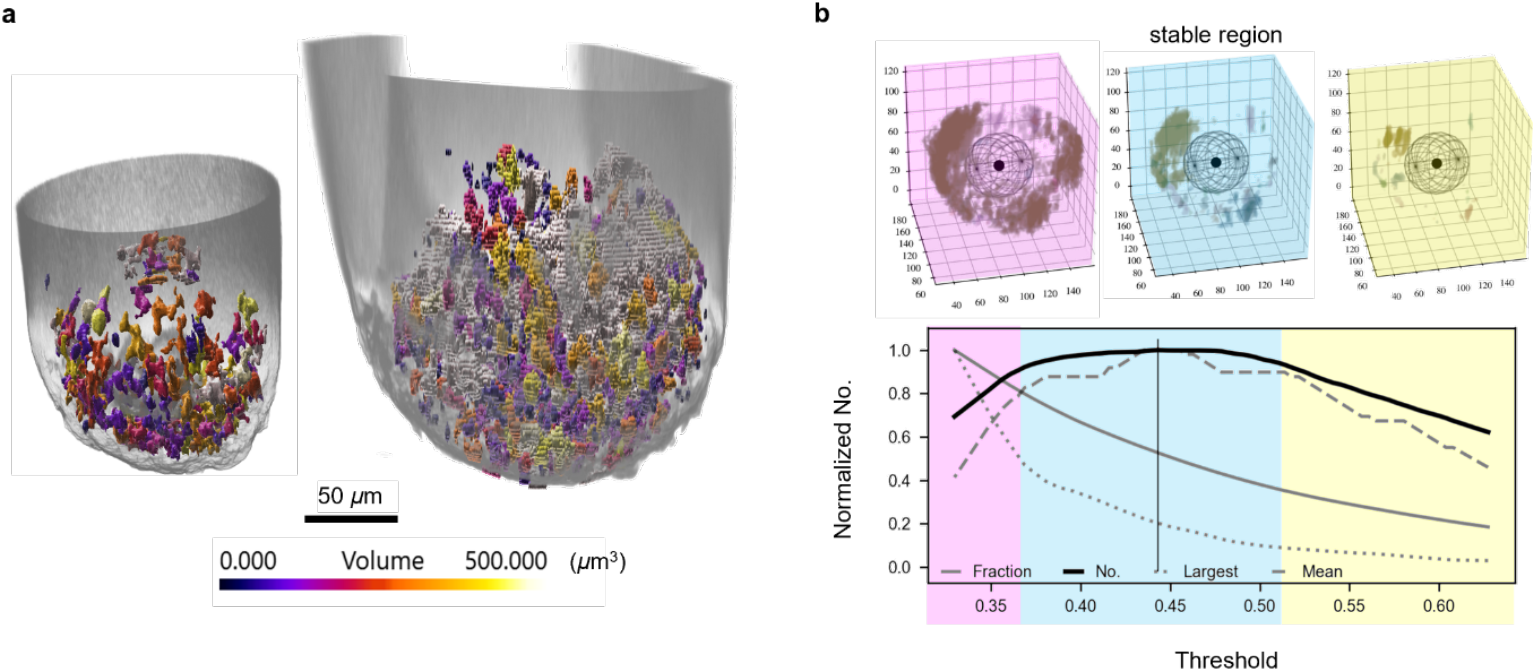
Characterisations of IG distribution and sensitivity test. **a**, Imaris 3D rendered IGs in a representative secondary (<180 µm, left) and tertiary (> 180 µm, right) follicle stained with 4kDa Dextran-FITC, using two-photon timelapse imaging. The IGs are colour coded according to their volume. **b**, Top: Same follicle with different thresholding conditions for the IGs. The different colours correspond to a lower threshold leading to over-detection of signals (pink), and a proper threshold exhibiting stable segmentation results (cyan) and a higher threshold leading to under-detection of signals (yellow). Bottom: Plot of IG properties against FITC intensity threshold: normalised IG volume fraction (grey solid line), normalised IG number (black solid line), normalised largest 3D rendered IG volume (grey dotted line) and normalised mean IG volume (grey dashed line).

**Extended Data Fig 2:**
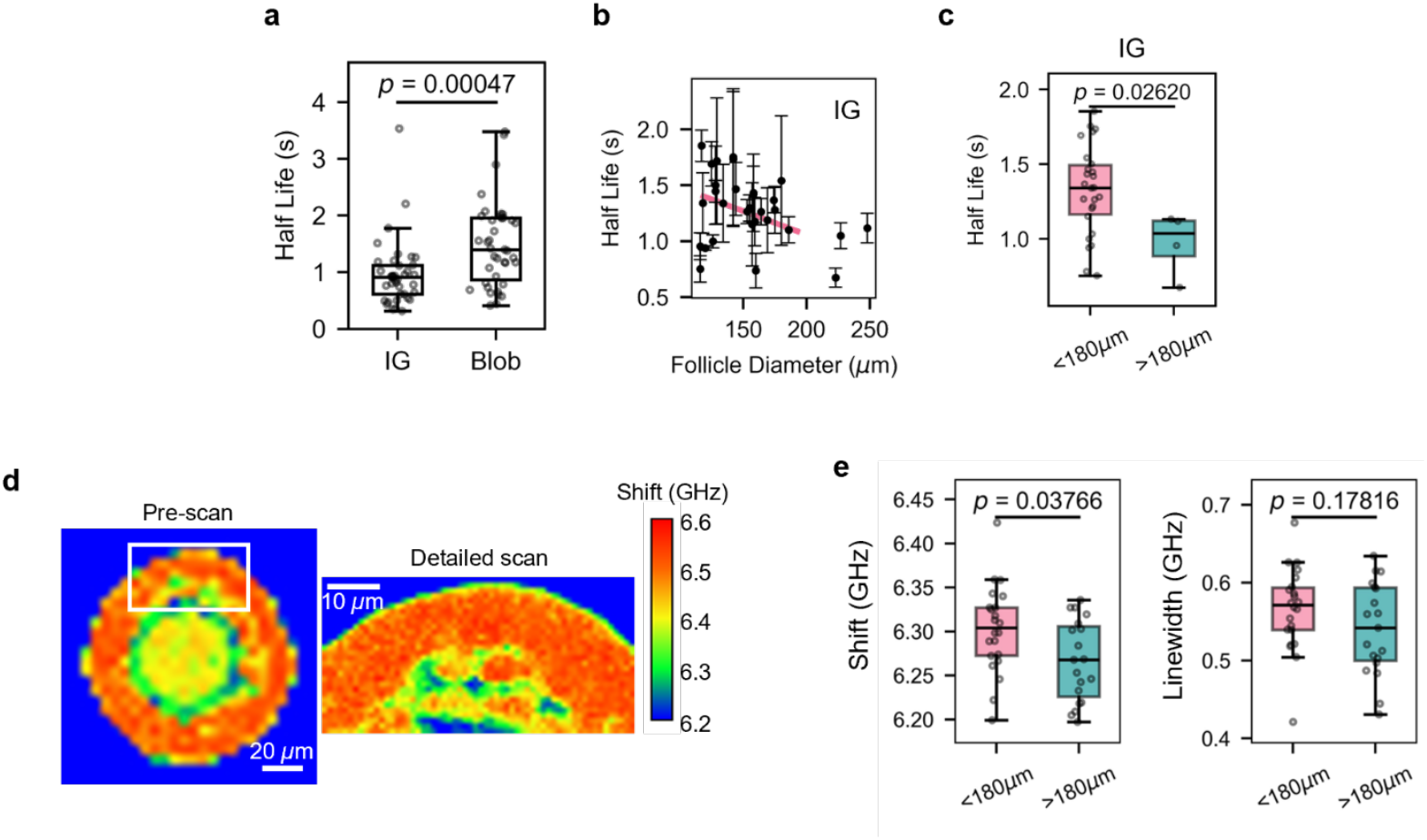
IGs are more viscous in secondary follicles. **a**, Boxplot of fluorescence recovery after photobleaching (FRAP) half-life in IGs and blobs (N = 4, n = 39 gaps from 14 follicles). **b**, Plot of average FRAP half-life of IGs in follicles against follicle diameter (Pearson correlation *R* = -0.312, p = 0.046). Red line demonstrates the trend of negative correlation. **c**, Boxplot of FRAP half-life of IGs in follicles smaller and larger than 180 µm. N = 2, n = 26 (small), 4 (large). **d**, Representative map of Brillouin shift for follicles (left). Inset shows zoomed-in scan of white box (see Methods). **e**, Boxplots of mean Brillouin shift (left) and linewidth (right) of IGs in isolated follicles smaller and larger than 180 µm. N = 4, n = 22 (small), 19 (large).

**Extended Data Fig 3:**
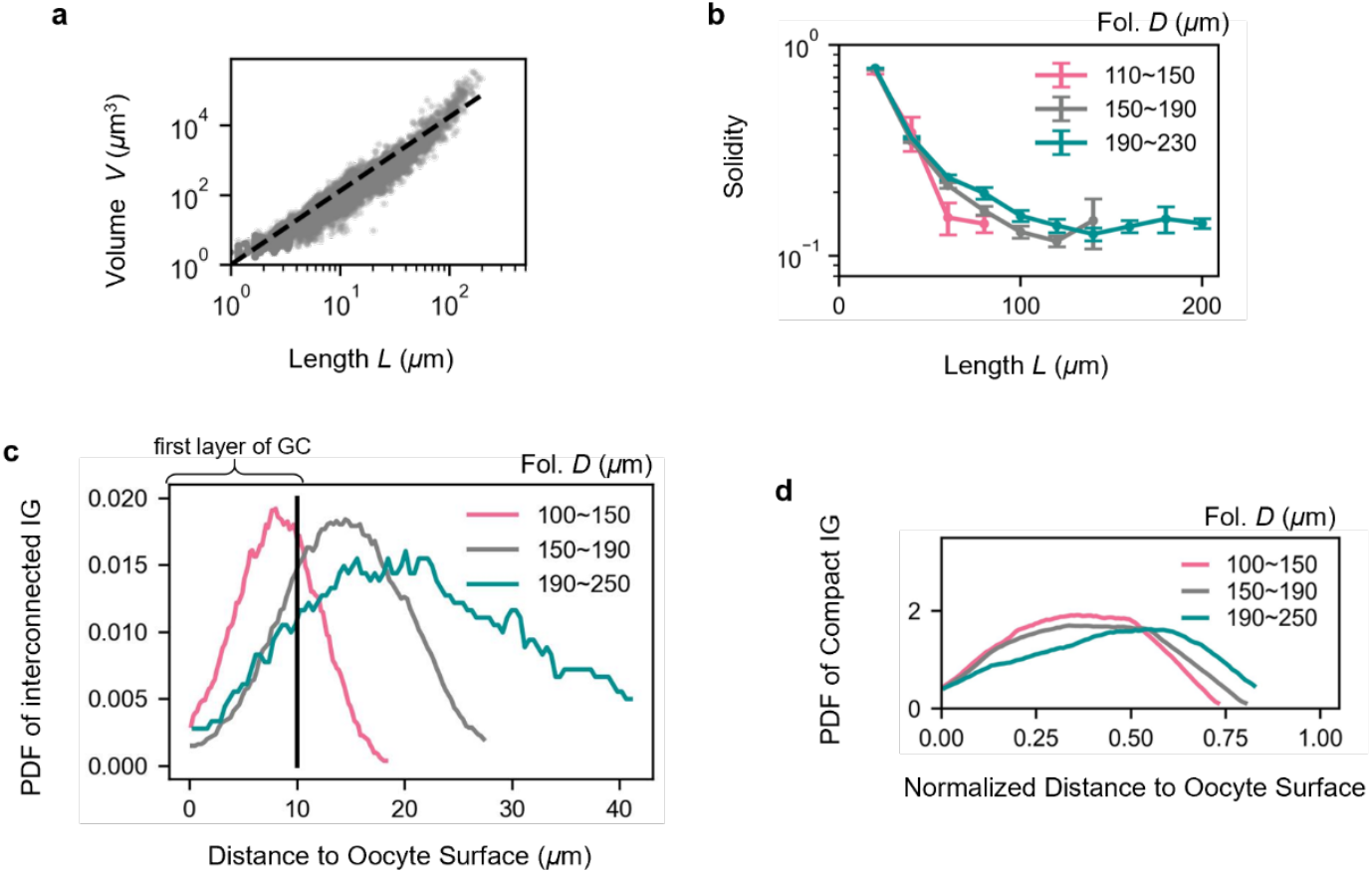
Further geometric analysis of IG morphology. **a**, Pooled data for plots of IG volume *V* against corresponding IG box length *L*. Dashed line indicates the power of log-log fitting, which is an overall IG effective dimension across all the samples. **b**, Pooled data for plots of IG solidity against its corresponding box length for different ranges of follicle sizes *D*. Error bar represents standard error of mean. **c**, Plot of probability density function (PDF) of IGs with solidity <0.25 over the absolute distance from oocyte surface, for different ranges of follicle sizes *D*. Black line corresponds to the first layer of GC. **d**, Plots of PDF of IGs with solidity >0.75 for different ranges of follicle sizes *D*. All data from N = 15, n=153 follicles.

**Extended Data Fig 4:**
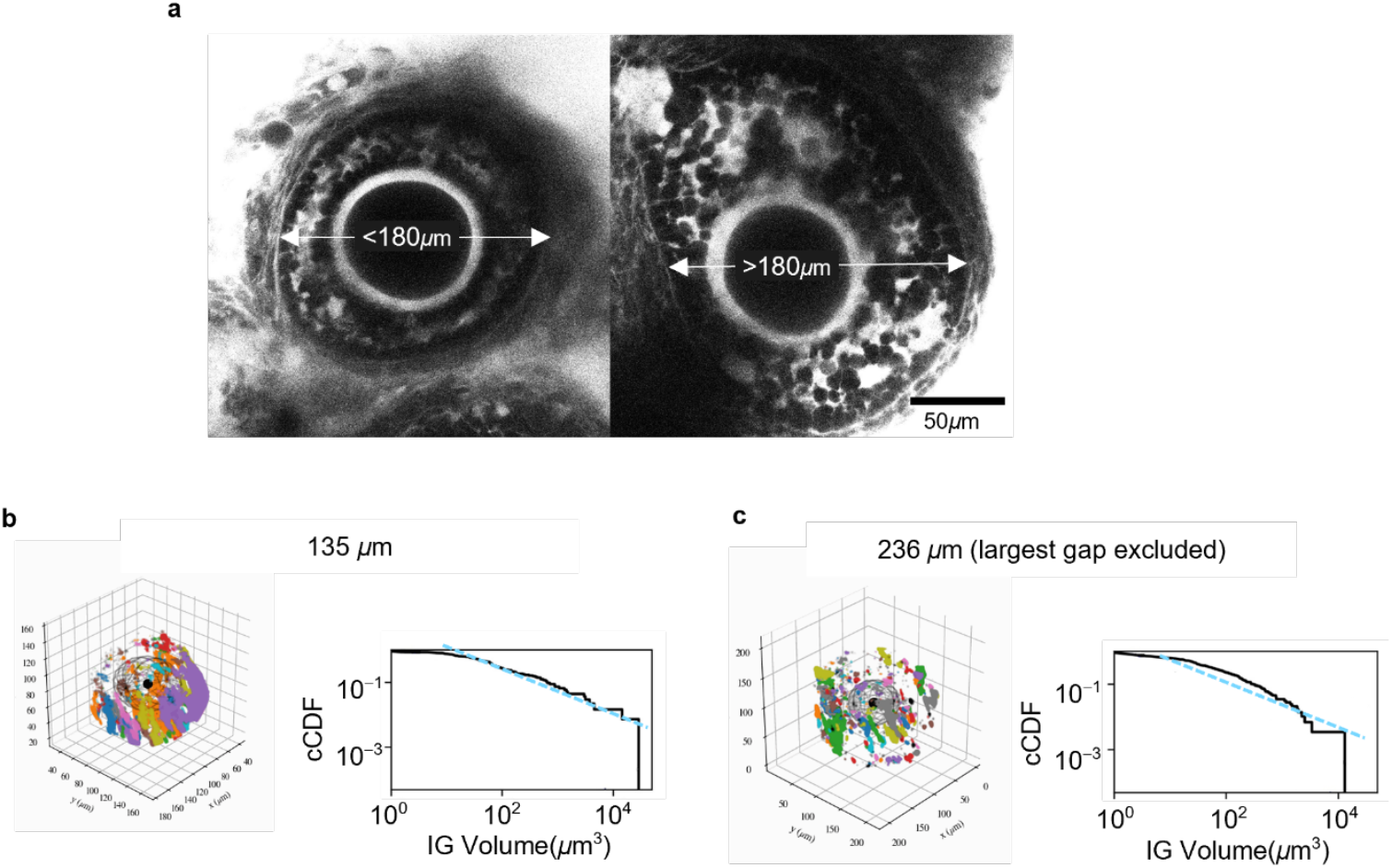
IG distribution of follicles in ovarian microtissues. **a**, Representative image of secondary (left) and tertiary (right) follicles in ovarian microtissue stained with 4kDa Dextran-FITC. **b**, 3D rendered IGs (left) and its corresponding cCDF plot (right) for a representative secondary follicle. **c**, 3D rendered IGs (left) and its corresponding cCDF plot (right) for a representative tertiary follicle. Colour codes indicate distinct IGs.

**Extended Data Fig 5:**
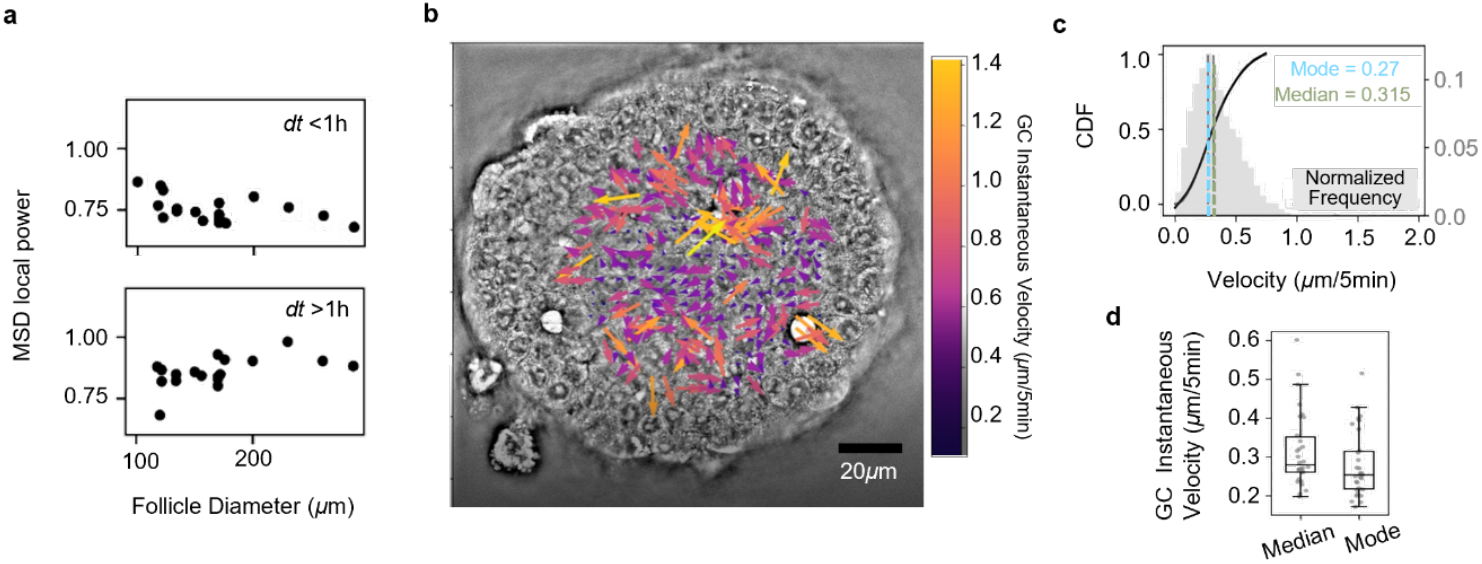
Characterisation of GC dynamics using quantitative phase imaging. **a**, Plot of MSD local power against follicle diameter for time intervals less (top) and more (bottom) than 1 h (N = 4, n =19). **b**, Representative image of a follicle imaged using holotomography. Velocity field from PIV analysis is overlayed and velocity is represented by the heat map. See Supplementary Video 6. **c**, CDF plot of GC velocity, extracted from **b**. GC median velocity is demarcated by the green dotted line while its mode is demarcated by the blue dotted line. **d**, Boxplot of the GC median and mode velocity (N = 5, n = 30).

**Extended Data Fig 6:**
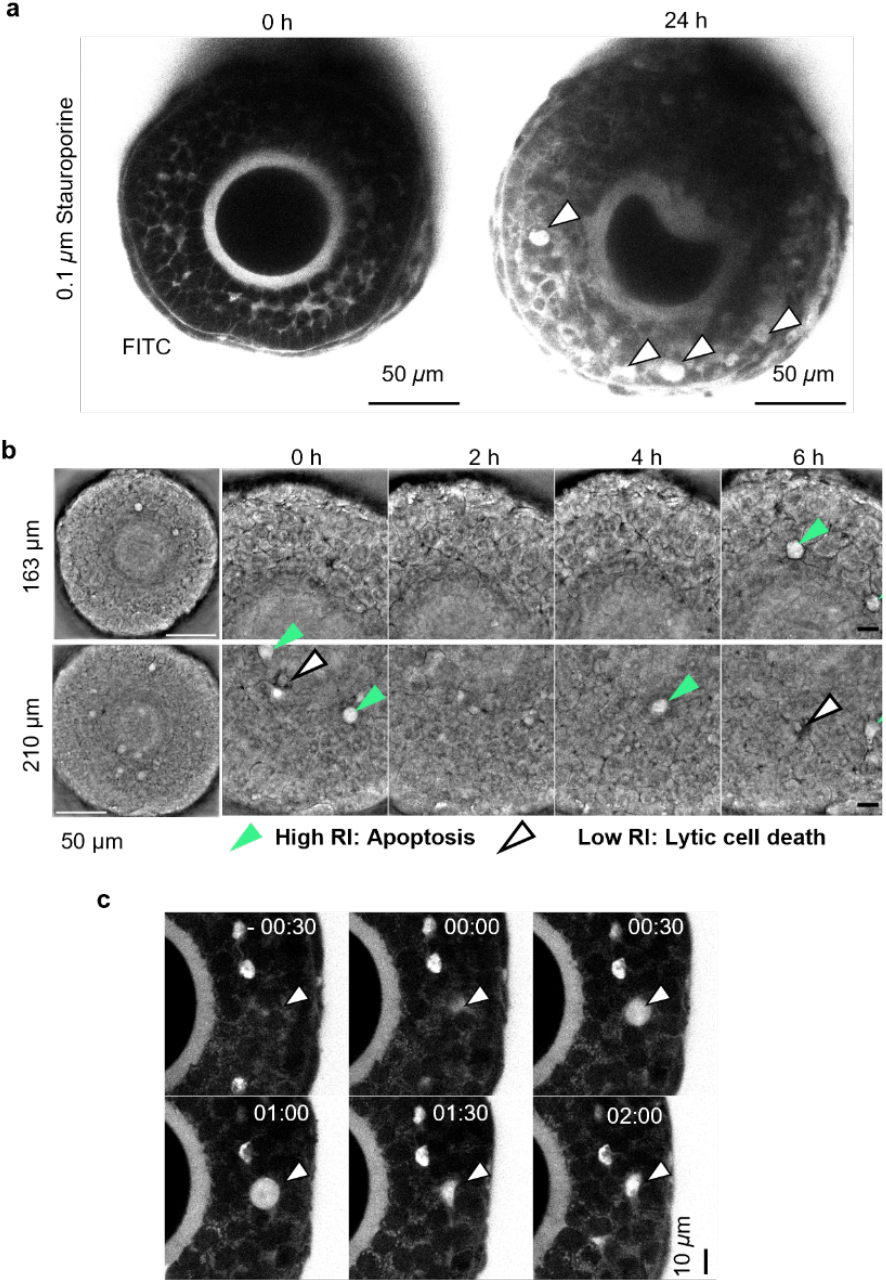
Emergence of non-apoptotic cell death in follicles larger than 180 µm. **a**, Representative images of follicles stained with FITC before (left) and after 24 h of DMSO (top) or staurosporine (bottom) treatment. White arrowheads indicate FITC blobs. **b**, Holotomographic images of isolated follicles at different time points, for follicles smaller (top) and larger (bottom) than 180 µm. Green arrowheads indicate high refractive index regions (apoptotic cells) while white arrowheads indicate low RI regions (cell death). See Supplementary Video 7. **c**, Representative images showing the emergence and disappearance of FITC blobs in FITC stained isolated follicle. Time is shown in hh:mm.

**Extended Data Fig 7:**
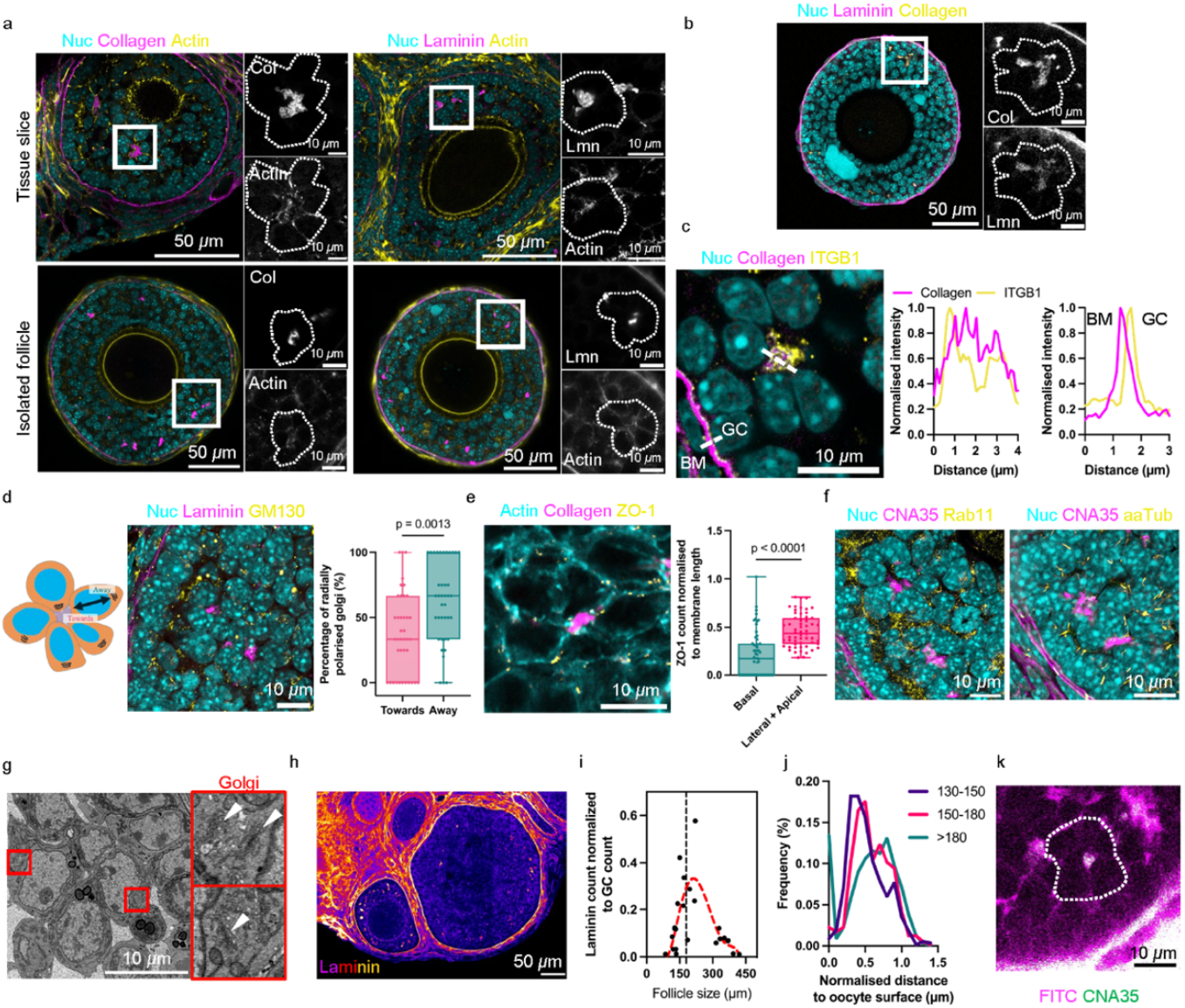
GCs exhibit ECM-enriched rosettes in secondary follicles. **a**, Representative images of follicles from ovary tissue slice (top) and isolated follicle (bottom) stained for nucleus (cyan), collagen (magenta, left) or laminin (magenta, right), and actin (yellow). Accompanying images to the right of follicles are the corresponding zoomed-in images from the white boxes. **b**, Representative image of a follicle co-stained with nucleus (cyan), collagen (yellow) and laminin (magenta). Inset: Zoomed-in image of white box, showing co-localisation of collagen and laminin in the rosettes. **c**, Left: Zoomed-in image of a follicle stained for nucleus (cyan), collagen (magenta) and ITGB1 (integrin beta-1, yellow). Right: Plots of line intensity for pan-collagen and ITGB1 across a GC rosette (left) or basement membrane (right). **d**, Illustration of defined Golgi orientation used for quantification. Representative image of a follicle in tissue slice, stained for nucleus (cyan), laminin (magenta) and Golgi apparatus (yellow). Right: Boxplot of percentage of radially polarised Golgi located towards or away from rosette (N = 4, n=11 follicles, 39 rosettes). **e**, Representative image of follicle from ovary tissue slice stained for nucleus (cyan), collagen (magenta) and ZO-1 (yellow). Right: Boxplot of ZO-1 count normalised to membrane length for basal and lateral+apical surface (N = 3, n = 23 follicles, 56 rosettes). **f**, Representative image of follicle from tissue slice stained for nucleus (cyan), collagen (magenta) and Rab11 (yellow, left) or acetylated alpha tubulin (yellow, right). **g**, Representative image of rosette in follicle from tissue slice imaged with SEM. Red boxes are zoomed in images of the Golgi apparatus (white arrowheads) of GCs in the rosettes. **h**, Representative image of follicles in ovarian tissue slice, showing rosettes stained with laminin (heatmap). **i**, Plot of laminin count normalised to the total number of GCs against follicle diameter (N = 4, n = 24). **j**, Probability distribution function of CNA35 count against normalised distance to oocyte surface (N = 3, n = 27). **k**, Representative image of rosette stained with FITC (IG, magenta) and CNA35 (pan-collagen, green) in isolated follicle. Dotted polygons represent the outline of rosettes.

**Extended Data Fig 8:**
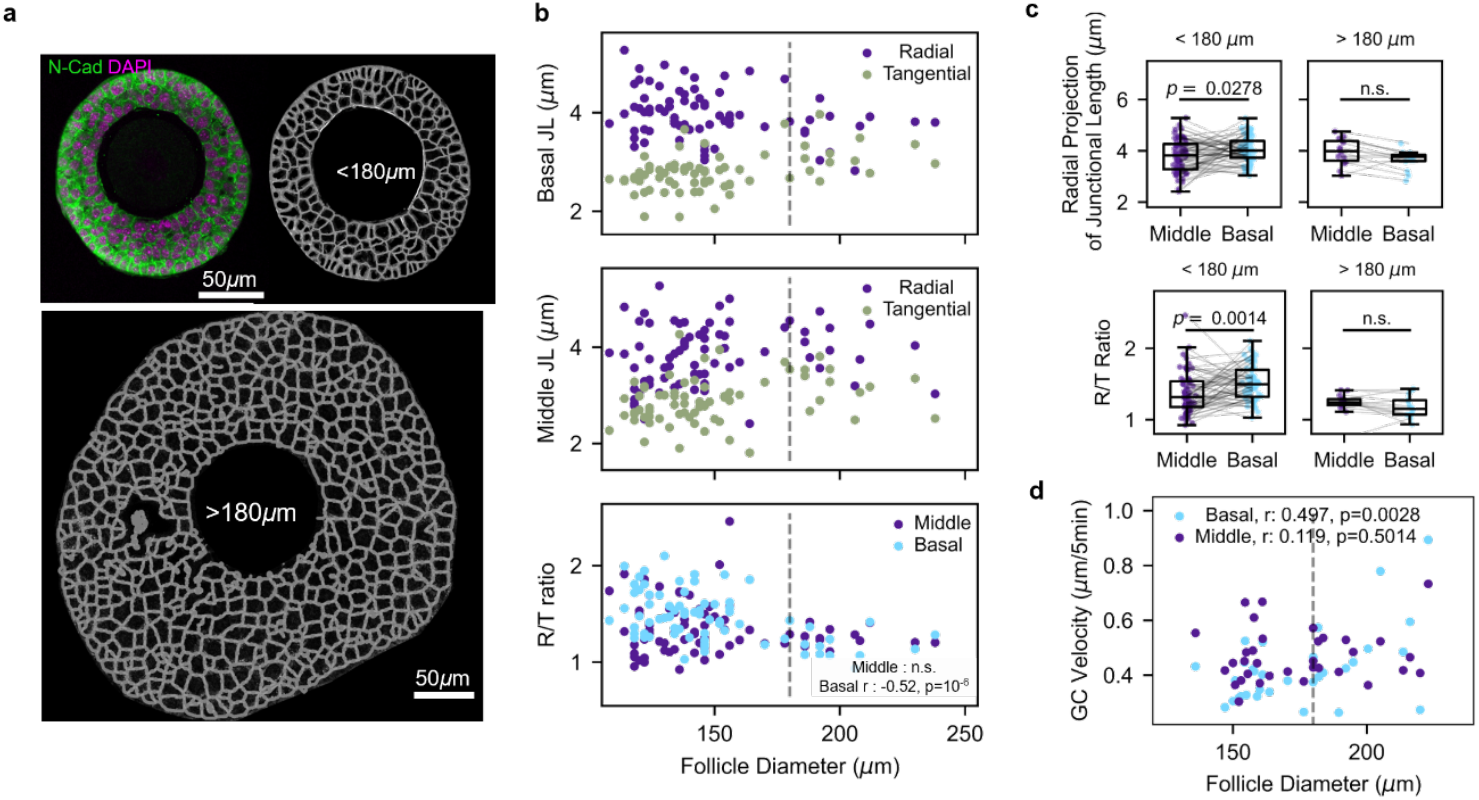
Characterisation of GC morphology and junctional properties in secondary and tertiary follicles. **a**, Representative image of N-cad and DAPI stained image of an isolated follicle (top left). N-cad intensity was used to segment GC cell boundaries for follicle smaller (top right) and larger than 180 µm (bottom). **b**, Plots of basal (top) and middle GC (middle) junctional length (JL) in the radial and tangential direction against follicle diameter. A ratio between the radial and tangential (R/T) JL for both the basal and middle GC were taken and plotted against follicle diameter (bottom). **c**, Pairwise boxplot of radial JL (top) and R/T ratio (bottom) for both middle and basal GC in the same follicle, for follicles smaller (left) and larger (right) than 180 µm. **d**, Plot of basal and middle GC velocity against follicle diameter. N = 2, n = 29 for **b** and **c**. N = 5, n = 40 for **d**. All dashed lines indicate 180 µm.

**Extended Data Fig 9:**
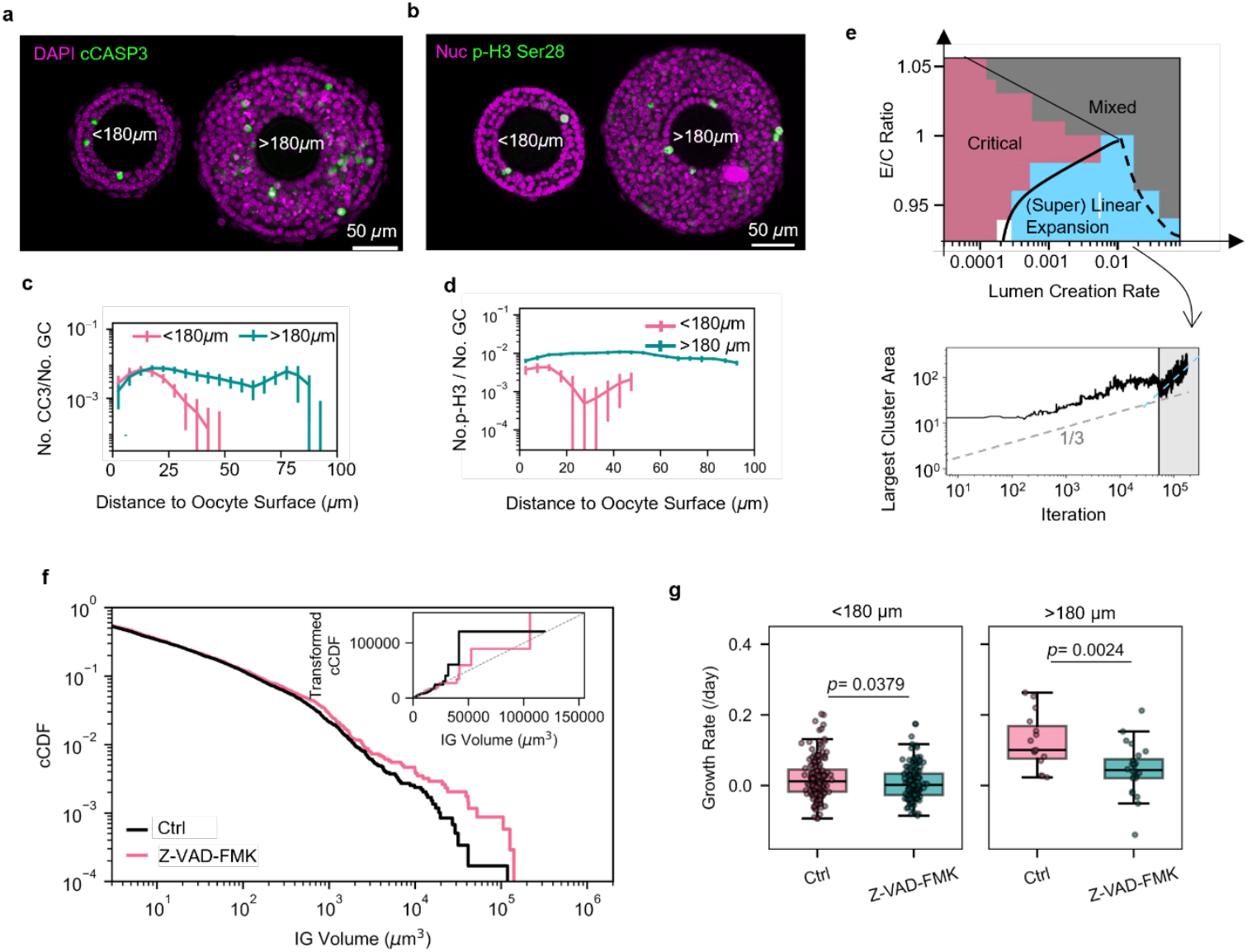
Non-conservative binary fluid model incorporating cell death and proliferation. **a-b**, Representative images of cCASP3 stained follicles (a) and p-H3Ser28 stained follicles (b). **c-d**, Plots of pooled average for normalised number of cCASP3 (**c**, N=4, n=55 (small), 37 (large)) and normalised number of p-H3 (**d**, N=6, n=49 (small), 50 (large)), against distance to the oocyte surface. Error bars represent S.E.M. **e**, Top: Phase diagram of 2D simulation with non-conservative gap-cell dynamics, starting from a dynamically critical state as the initial state. Gap turnover rate is the number of gap creation event at a single step; E/C ratio is the number of gap elimination over gap creation at each time step. Bottom: Typical plot of time evolution of largest cluster in the phase of super linear expansion after non-conservative dynamics is implemented at 50000th iteration. **f**, Plots comparing pooled cCDF for IG volumes in isolated follicles larger than 150 μm between control (DMSO) and pan-apoptosis treatment (Z-VAD-FMK), 3 days after treatment (N = 3, n = 14 (DMSO), 10 (Z-VAD-FMK) follicles); inset: transformed cCDF against IG volume on linear-linear scales. **g**, Boxplots for growth rate of for control (DMSO) and Z-VAD-FMK-treated follicles smaller (left, N=5, n=163 (ctrl), 142 (Z-VAD-FMK)) and larger than 180 μm (right, N=5 , n=15 (ctrl), 24 (Z-VAD-FMK)).

**Extended Data Fig 10:**
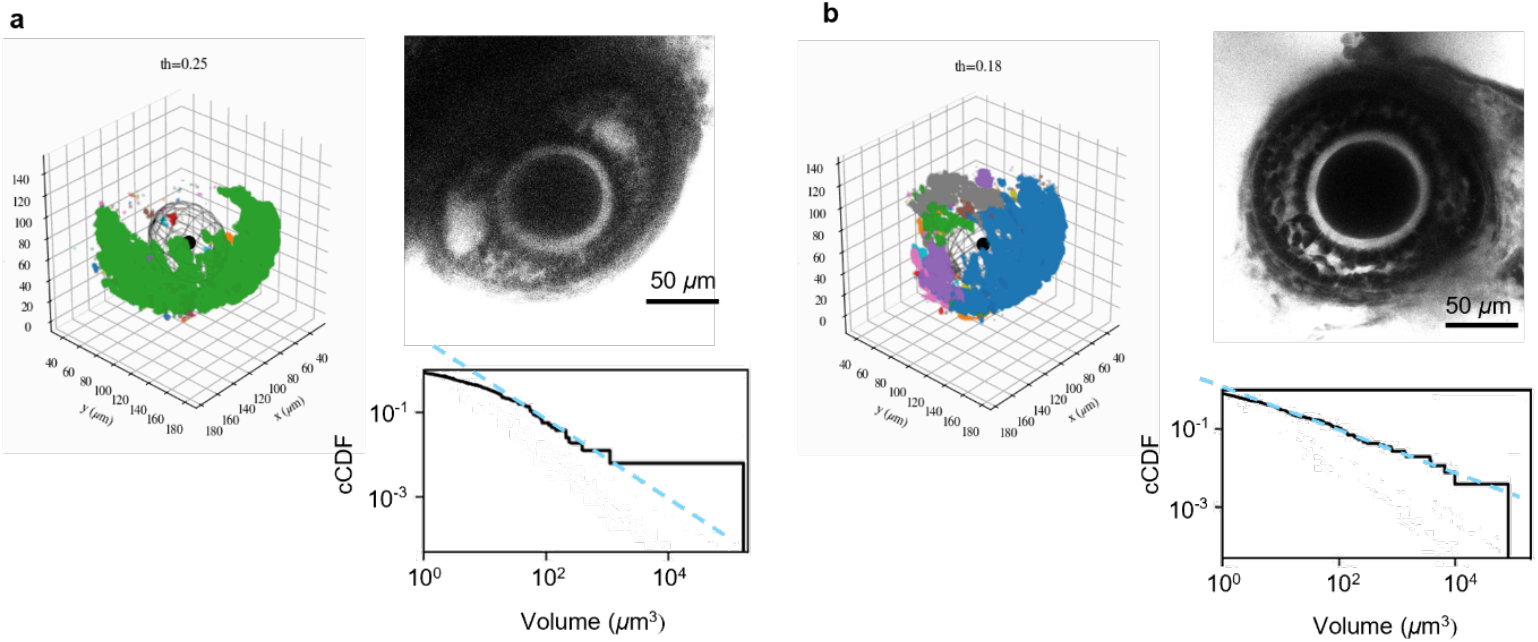
Deformed follicles *in situ* appear to exhibit spatially resolved fluid gaps. **a**, 3D rendered IGs in deformed follicle (left), which is stained with 4 kDa Dextran-FITC in microtissue (top right). Corresponding cCDF plot of IG volume showed a poor fit to power law behaviour (bottom right). **b**, 3D rendered IGs in a non-deformed follicle which is similar in size to that of **a** (left). FITC staining revealed more percolative IGs (top right), with corresponding cCDF plot of IGs showing a better fit to power law behaviour (bottom right). Color codes represent distinct IGs.

## METHODS

### Animals

C57BL/6NTac female mice, aged P25-P28, used in the experiments were group housed in individually ventilated cages with access to water and food under a 12-hr light/12-hr dark cycle. Mouse rooms were maintained at 18-25 °C and 30-70% relative humidity. The mice were euthanized by carbon dioxide asphyxiation followed by cervical dislocation. All mice care and use were approved by the Institutional Animal Care and Use Committee (IACUC) at the National University of Singapore.

C57BL/6NTac female mice, aged P21, used in Brillouin imaging (Extended Data Fig. 2d) and R26-H2B-mCherry female mice, aged 48 weeks, used in timelapse imaging (Fig. 3a, b) were obtained after they were euthanized by cervical dislocation from the animal facility at the European Molecular Biology Laboratory, with permission from the institutional veterinarian overseeing the operation (ARC license number 21-008_HD_LAR). The animal facilities are operated according to international animal welfare rules (Federation for Laboratory Animal Science Associations guidelines and recommendations). Ovaries were dissected from the mice and transferred to an isolation buffer consisting of Leibovitz’s L15 medium (ThermoFisher, 21083027) supplemented with 0.3% Bovine Serum Albumin (BSA) (Sigma-Aldrich, A9418).

### 3D follicle culture

Follicles were mechanically isolated from dissected ovaries under the stereomicroscope attached to thermal plate using tweezers in isolation buffer at 37 °C. Growth medium, consisting of MEM-α GlutaMAX (ThermoFisher, 32561037) supplemented with 10% Fetal Bovine Serum (ThermoFisher, 10082-147), 1% Penicillin-Streptomycin (ThermoFisher, 15140-122), 1X Insulin-Transferrin-Selenium (Thermo Fisher, 41400045), and 50 mIU/ml follicle stimulating hormone (Sigma-Aldrich, F4021-.1MG) was prepared. Individual follicles were transferred to an agarose microwell and cultured in growth medium at 37 °C, 5% CO_2_, 95% humidity overnight. 1% alginate (Sigma-Aldrich, 71238) was prepared in phosphate buffer saline (ThermoFisher, 18912014) and mixed with growth medium in a 1:1 ratio. Follicles were submerged in the 0.5% sodium alginate solution, and hydrogels were formed by mouth pipetting crosslinking solution consisting of 50 mM calcium chloride and 140 mM sodium chloride into individual microwells. The alginate was allowed to crosslink for about 1 min. For long term cultures, half the volume of the growth medium was changed in each well every two days.

### Microtissue culture

Microtissue culturing protocol was adopted from. In brief, RGD-modified dextran hydrogels were prepared using TrueGel3D Hydrogel Kit (Sigma-Aldrich, TRUE7) according to manufacturer’s instruction. After addition of crosslinker, 3 µL of RGD-modified dextran hydrogel were pipetted onto a 30 mm PTFE cell culture insert (Merck, PICM0RG50) to form hydrogel domes. The hydrogels were incubated at room temperature for 30 min before the addition of 1X PBS followed by incubation at 37 °C for 1 h. Prior to the completion of incubation, live microtissues were prepared by slicing up ovaries into small chunks of 5-10 follicles using a 27-gauge needle. After the incubation, the 1X PBS was aspirated and the microtissues were placed on top of the hydrogel domes. 1.4 mL of growth media was added to the well of a 6-well culture plate and the cell culture insert was placed in the well. 5 µL of growth media was pipetted over the microtissue-hydrogel assembly to wet the area and ensure a thin film of media dome for an air-liquid interface culture. Half the volume of the growth medium was changed in each well every two days. The microtissue-hydrogel assembly was wetted following every medium change.

### Pharmacological treatments

For isolated follicles cultured in 3D, the day of alginate embedment was dictated as Day 0. Drug were treated on Day 0. Microtissues were cultured for 2 days for the microtissue to attach to the RGD-modified dextran hydrogel. Drug were treated on Day 2 during the medium change. ADH-1 (Selleckchem, P1239) was used at 200 µM in DMSO and 1 mM in PBS to inhibit N-cadherin cell adhesion. Staurosporine (Sigma-Aldrich, S5921) was used at 100 nM to induce cell death. Z-VAD-FMK (Selleckchem, S7023) was used at 80 µM to inhibit apoptosis.

### Tissue sectioning

Ovaries were fixed in 4% paraformaldehyde (Santa Cruz Biotechnology, sc-281692) at room temperature for an hour. For vibratome sectioning, the fixed ovaries were embedded into 4% low-melting point agarose (ThermoFisher, 16520050). The embedded ovaries were sliced into 150 µm thick tissue sections using a vibratome (Leica, VT1200) in PBS at 0.05 mm/s speed and 1 mm amplitude. For cryosectioning, the fixed ovaries were equilibrated in 15% and 30% sucrose solution in 1X PBS for 1 h sequentially followed by FSC 22 Frozen Section Media (Leica, 3801480) and snap frozen using dry ice before they are kept frozen at –80 °C. The cryopreserved ovaries were sliced into 15 µm thick tissue sections using a cryostat (Leica, CM1950). Live microtissues were prepared by slicing up ovaries into small chunks of 5-10 follicles using a 27-gauge needle.

### Immunofluorescence staining

Isolated follicles were fixed in 4% PFA at room temperature for 30 min and washed with washing buffer thrice before immunostaining. Fixed samples were incubated in blocking-permeabilizing solution (3% BSA and 0.3% Triton X-100 in 1X PBS) at room temperature for 2-4 h. If mouse antibody is used, an additional blocking step using M.O.M.® (Mouse on Mouse) Blocking Reagent (Vector Laboratories, MKB-2213-1) was carried out according to manufacturer’s protocol before resuming primary antibody staining. Permeabilised and blocked samples were incubated at 4 °C in primary antibodies diluted in blocking solution overnight. The tissues were washed 5 times in washing buffer before they were incubated in secondary antibodies diluted in the washing buffer for 1-2 h at room temperature. They were washed thrice in washing buffer before mounting. Rosette staining using rabbit anti-laminin conjugated to PE/Atto594 (1:100, Novus Biologicals, NB300-144PEATT594) was carried out after secondary antibody staining at 4 °C in blocking solution overnight. After which, the tissues were washed thrice in washing buffer before mounting. The ovarian slices were mounted using ProLong Gold antifade mountant (ThermoFisher, P36930) and left to cure overnight at room temperature, whereas isolated follicles were mounted using SlowFade Gold antifade mountant (ThermoFisher, S36936) prior to imaging.

Primary antibodies used were rabbit anti-cleaved caspase 3 (1:100, Abcam, ab2302), rabbit anti-GM130 (1:200, abcam, ab52649), rabbit anti-N-cadherin (1:300, Proteintech, 22018-1-AP), rabbit anti-ZO-1 (1:100, Thermofisher, 61-7300), rabbit anti-Rab11 (1:100, CST, 5589S) rat anti-integrin β1 chain (1:100, BD Biosciences, 553715), mouse anti-acetylated alpha tubulin (1:100, Sigma-aldrich, T-7451),. Alexa Fluor 488 labelled anti-mouse (1:300, Invitrogen, A11055) and Alexa Fluor 633 labelled anti-rabbit (1:300, Invitrogen, A21070) was used as secondary antibodies. DNA was stained with DAPI (2 µg/mL, Sigma-Aldrich, D9542) and F-actin was stained with either Alexa Flour 488-labelled phalloidin (1:300, Invitrogen, A12379) or Alexa Flour 633-labelled phalloidin (1:300, Invitrogen, A22284). Collagen was stained with either CNA35-EGFP or CNA35-mCherry at 8 µM.

Fixed isolated follicles and tissue sections were imaged on Nikon A1Rsi confocal microscope with NIS Elements using Apo 40×/1.25 WI Lambda S DIC N2 objective or at a higher magnification using Plan Apo 100x/0.13 objective at 1-2 µm z-slices. Zeiss LSM980 confocal microscope was also used for imaging, and operated with ZEN Blue using Plan-Apochromat 20x/0.8. Higher magnification images were captured using Plan-Apochromat 63x/1.4 Oil DIC M27 at 2 µm z-slices. Stitching was carried out with 10% overlap using lasers 405 nm, 488 nm, 561 nm, 640 nm.

### Live imaging

Follicles cultured in 3D and stained with FITC at 160 µM were used for IG imaging on Zeiss LSM980 for IG distribution analysis. Experiments were carried out on the 40x lens at 2 µm z step. Control follicles were imaged before 5 h of drug treatment followed by another round of imaging post drug treatment. Microtissue stained with 4kDa Dextran-FITC at 400 µM were used for IG imaging on Zeiss LSM980 for IG distribution analysis. The experiments were carried out on the 40x lens at 2 µm z step.

For temporal correlation analysis, follicles cultured in 3D and stained with FITC at 160 µM were used for timelapse imaging on Zeiss LSM980. Experiments were carried out on the 40x lens at 1 µm z step and 5 min time interval for a total of 6 h duration. Control follicles were imaged for 3 h before drugs were added and imaged for another 3 h. Follicles cultured in 3D and stained with 4kDa Dextran-FITC at 400 µM were used for timelapse imaging on Zeiss LSM980 for tracking follicle growth. The experiments were carried out on the 20x lens with 1.8-2x zoom at 2.5 µm z step and 10 min time interval for up to 80 h. Control and ADH-1 treated follicles were imaged separately. H2B-mCherry follicles were used for timelapse imaging on Zeiss LSM880 for MSD tracking. The experiments were carried out on the 40x lens at 2 µm z step and 10 min time interval for a total of 16 h duration. Control and ADH-1 treated follicles were imaged separately.

Live follicles in 3D culture were imaged on Zeiss LSM980 on the non-descanned two-photon mode at 800 nm wavelength using LD C-Apochromat 40x/1.1 W Corr M27. H2B-mCherry live imaging were carried out on Zeiss LSM880 on the non-descanned two-photon mode at 1080 nm wavelength using LD C-Apochromat 40x/1.1 W Corr M27 at 2 µm z step and 10 min time interval for a total duration of 16 h.

### Fluorescence recovery after photobleaching (FRAP)

FRAP experiment on IGs and blobs was carried out on 4kDa Dextran-FITC stained follicle using Zeiss LSM980. The experiment was set up using the FRAP module in ZEN Blue software such that images were acquired continuously at about 0.35 s per frame for 15 frames before bleaching at 100% power using a 488 nm laser at a scanning speed of 8 over two 2 µm by 4 µm regions. Postbleaching recovery was taken for 30 frames. This was repeated over four iterations in a total of 120 frames. An additional control region without bleaching was also included to correct for any photobleaching.

FRAP experiment on IGs in varying follicle sizes was carried out on 10 kDa Dextran-FITC stained follicle using Nikon TiE fitted with Yokogawa CSU-W1, Gataca iLas Ablation system and Photometrics Prime 95b CMOS camera. The experiment was set up using the Gataca module on MetaMorph such that images were acquired continuously at about 0.07-0.08 s per frame for 150-200 frames and a pre-set 3.5 µm diameter circle region was bleached manually at 10% power using a 488 nm laser for 0.5 s. Postbleaching recovery was taken for about 5 s before a second round of bleaching.

Intensity and timepoints were exported for processing. Intensities in the ROI were first subtracted by the intensities in the control region at the corresponding timepoint to correct for any photobleaching. Postbleaching recovery was normalised to the average intensity 3-5 s prior to bleaching. The processed data were plotted, and curve fitted using a non-linear one-phase association equation to obtain the halftime and plateau in GraphPad Prism 9. Diffusion rate *D* is estimated, in literature^34^, as 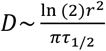, where *r* is the effective radius of bleaching region (∼3 µm), and τ_1/2_ being the half-life obtained from the FRAP experiment.

### Brillouin microscopy

Brillouin imaging was performed using a modified version of the Brillouin microscope previously described in^13^. Briefly, this consists of a commercial Zeiss body (Axiovert 200 M) coupled with a home-built spectrometer based on a 2-VIPA configuration and a Lyot stop. The theoretical spectral resolution of 270 MHz is subtracted from the measured linewidth values to deconvolve them, in the assumption of Lorentzian lineshape. A 660nm laser (Torus, Laser Quantum) is used for Brillouin imaging. Brillouin images are acquired with a 40×1.0NA Zeiss objective and an integration time for a single point of 100 ms, while keeping the optical power on the sample below 6 mW. A 488 nm laser, coaligned with the 660 nm laser, allows for confocal imaging of GFP fluorophores.

Live follicles cultured in agarose microwells were used for imaging. A pre-scan using 5 µm pixel resolution was carried out to locate IGs before an actual scan of 1 µm pixel resolution at the IGs to reduce imaging time. IGs were segmented using the Brillouin shift channel and was independently checked for co-localisation with FITC staining imaged on the 488 nm laser to ensure faithful segmentation of IGs. A binary dilation of 1 µm and subsequent erosion of 3 µm on the segmented areas were carried out to avoid GC-IG boundaries. Brillouin shift and linewidth were extracted for each pixel and averaged for each IG before they were then averaged for each follicle. Datapoints with linewidth fitting, R2, greater than 0.92 were taken.

### Quantitative phase microscopy

Holotomograms of 3D cultured follicles were obtained using a low-coherence HT system (HT-X1, Tomocube Inc., Korea). To ensure the maximal depth of view coverage on the HT-X1 system, follicles were cultured in 3D in a customised PDMS U-shaped microwell with 2 mm diameter and 2 mm depth. The microwell mold was designed and 3D printed using clear resin V4 on Form 3 3D printer (Formlabs). A 25 µm layer thickness was chosen for the print. After the print, the resin mold was soaked in IPA for 12 h before a fresh IPA rinse and was dried using nitrogen gas. The resin mold was then cured using Form cure (1st generation) for 15 min at 60 °C. A 1:10 curing agent to base ratio of Sylgard™ 184 Silicone Elastomer (PDMS) was used to cast the microwell from the resin mold. The PDMS was degassed for 10 min before curing for 4 h in an 80 °C oven. A 1 mm dermal punch was used to perforate the bottom of the wells before the molded PDMS was placed on a bioinert 35 mm μ-dish (ibidi, 81150). To encapsulate the follicles, the wells were first filled with 4 μL of 0.5% sodium alginate solution before follicle were submerged in the wells. Another 3 μL of crosslinking solution was then added and incubated for 2 min for the alginate gel to form. 2 mL of growth medium was added to the dish for the 3D culture. For long term cultures, half the volume of the growth medium was changed in each dish every two days.

HTX1 employs incoherent 450-nm LED light for illumination and follicles were imaged at a z step of 0.78 µm over a range of about 142 μm. For PIV measurements, timelapse were captured at 5 min time intervals up to 10 h. Control follicles were imaged for 3-5 h before drugs were added and imaged for another 3-5 h. Captured data were used to reconstruct the RI of the samples within the desired field of view in the HTX processing software. All motorized microscopic operations were controlled and monitored by an operating software TomoStudio X (Tomocube Inc., Korea).

### Electron microscopy

Ovaries were fixed with 3% PFA and 2% glutaraldehyde (GA) overnight at 4°C. They were washed thrice with PBS for 5 min each. The ovaries were either processed as a whole organ or 150 μm tissue slices using the vibratome. The samples were incubated in 1% osmium tetraoxide (OsO4) with 1.5% potassium ferrocyanide in PBS for 1 h on ice and then washed thrice with distilled water for 5 min each. The samples were then placed into 1% thiocarbohydrazide (TCh) in distilled water for 20 min at room temperature and washed thrice with distilled water for 5 min each. The samples were then placed into 1% OsO4 in distilled water for 30 min at room temperature and washed thrice with distilled water for 5 min each. They were next incubated with 1% uranyl acetate (UA) in distilled water overnight at 4°C and washed thrice with distilled water for 5 min each. 0.02 M lead nitrate and 0.03 M aspartic acid were mixed well together, and the pH was adjusted to 5.5. The samples were kept in lead aspartate solution for 30 min at 60°C in the oven, and again washed thrice with distilled water for 5 min each. Tissues were dehydrated with ethanol, increasing gradually from 25%, 50%, 75%, 95% and 100%, with 10 min in each solution on ice before washing with acetone twice for 10 min each on ice. For resin infiltration, the samples were placed in a 1:1 acetone-araldite resin mixture for 30 min and then 1:6 mixture overnight. They were then transferred to pure araldite for 1 h in a 45°C oven. This was done thrice before they were transferred into an embedding mould with pure araldite and cured for 24 h at 60°C. The embedded samples were then sectioned using a Diatome diamond knife with the Leica UC6 ultramicrotome, and 100 nm ultrathin sections were collected onto silicon wafers. SEM imaging was done with Thermofisher FEI Quanta 650 FEG-SEM, where large area montage scans were acquired with MAPS 2.1 software using the backscatter mode (vCD detector) at 5 kV, 5 mm working distance.

### GC cell tracking

Follicles from F26-H2B-mCherry mouse were cultured in 3D and imaged with a time interval of 10 min for a total duration of 16h. The position of each nucleus *x*_*i*_ in 3D was detected and tracked using trackPy package. Mean square displacement (MSD) as a function of time interval Δ*t* was calculated as MSD= <(*x*_*i*_(*t*+Δ*t*)-*x*_*i*_(*t*))^2^>_*i,t*_, where the bracket average <·> is taken over all cells and all existing time points. A linear relationship MSD =*η*Δ*t* indicates a diffusive manner of cell movement as particles in the liquids. The diffusion coefficient is *η*/6 in 3D.

### FITC gap detection

3D stacks of FITC signals were pre-processed to similar levels of contrast and brightness across stacks using tophat and autolevel functions in Python package skimage. Oocyte and the basal GC layer were detected with canny algorithm and then removed from the original images. A proper intensity threshold was set to transform the images into binary values for FITC positive and negative pixels respectively. 3D agglomerative clustering algorithm with neighbour distance 1 was applied to segment FITC positive pixels into disconnected pieces of clusters. By scanning a large range of intensity thresholds, the optimal threshold was found so that the number of detected number of clusters is maximized. The information of segmented clusters of points under the optimal threshold were then passed to the following analysis of gap geometry and size. Blobs are detected by such criteria that 90 percentile of intensity in the pixel is larger than 60% of the maximum FITC intensity and the sphericity, ratio of square root of surface area over cubic root of volume normalized by 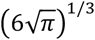 larger than 0.75.

### Quantification of N-cad junctional continuity

5-10 cell-cell junctions in the basal GCs were selected on the equatorial plane of the follicle using a 2-pixel wide line in the actin channel. The determined threshold was applied across the dataset to mask for N-cad signals in the junctions in both control and treated follicles. The same was carried out independently in the actin channel. N-cad junctional continuity in the basal GCs was measured as a fraction of the total area of masked N-cad signal over the total area of masked actin signal in the same cellular junction. An average was taken for each follicle. The same cell-cell junction selection was used to extract the N-cad fluorescence intensity. An additional normalisation to the background intensity in the nuclear region of the GC was carried out.

### Quantification of basal and mid GC velocity

The instantaneous speed of GC was determined by performing PIV measurements on 2D slices from holotomograms of 3D cultured follicles. A custom PIV analysis pipeline was implemented in Python using the Open PIV library. Image sequences were stacked and pre-processed with ImageJ macros for the detection of GC texture patterns. For each follicle, basal GCs were first identified, before mid GCs were identified at a plane 11.7 µm above the basal GC slice. Stable and non-rotating frames were selected for analyses, and a bandpass filter range of 0.00155 µm to 6.22 µm was applied to remove background noise. The PIV interrogation and search window sizes were set to 8.55 µm and 9.33 µm, respectively, corresponding to the typical diameter and displacement range of mid GCs (Extended Data Fig. 5b, left). Global displacement was removed by subtraction of global average of velocities. Outliers with a velocity magnitude above 0.5 µm/min or with a signal-to-noise ratio lower than 1 were removed before obtaining the median magnitude of velocity. Mode was obtained by taking the zero of second derivative of cumulative distribution function (CDF) of the velocity magnitude (Extended Data Fig. 5b, right).

### Quantification of T1 transition frequency

The basal GCs in a single 2D slice from holotomograms of 3D cultured follicles was manually tracked for T1 transition events over 2 h. A T1 event is classified by the loss of contact between two adjacent cells, and the subsequent establishment of contact between two new cells. T1 transition frequency per hour was determined by the number of T1 occurrences as a fraction of the initial number of basal GCs followed in the same section.

### IG geometrical dimension analysis

After gap detection algorithm was applied and blobs removed, clusters of distinct IGs were obtained for each follicle. Volume of IGs was quantified by counting the number of pixels in 3D and box length of IG was quantified by the largest distance between pairs of pixels in this cluster. The volumes of IGs grow with their box length and the effective dimension of IGs for each follicle was probed by fitting the linear coefficient of volumes against box lengths in the log-log scale.

### Solidity analysis

For each gap, the solidity is defined as the ratio between its volume and the volume of its convex hull. It is an 3D extension of the solidity used as shape descriptor for 2D objects^15,35–37^ As a ratio of real volume over the volume of convex hull, solidity measure how densely an object occupies the space. There is an exponential decay of solidity *S* with the IG length *L* to a finite residual (Extended Data Fig. 3c). From the decay *S*=exp(-*L*/*l*) we extracted the scale *l* for IG, ignoring the finite residue of low *S* at the large *L*. Then a characteristic length of decay *l*, named as solidity scale in right panel of Fig. 2c, was fitted from the decay for each follicle to indicate the typical IG size with which gaps are considered as solid.

### Temporal correlation analysis of FITC images

FITC timelapses over 5h with 5min interval of isolated follicles without rotational displacement were passed to image pre-processing using tophat functions to equalise the contrast across the space. The translational displacement in 3D was then removed by matching follicle boundary over time. We used the same gap detection method as elaborated above to segment FITC positive parts. The region of interest (ROI) was selected as part occupied by GCs between oocyte surface and basement membrane. All the pixels in ROI with FITC positive signals were assigned as 1 and other pixels in ROI as 0. The autocorrelation *C* as a function of time interval *dt* is calculated as

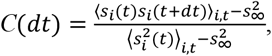

where *s* is the assigned value for FITC signals, the bracket <·> is taking average over all the pixel in the ROI and all time point and *s*_∞_ is supposed to be the steady state density of FITC positive signals. Practically we used the IG fraction for *s*_∞._ The correlation time τ was extracted by fitting correlation decay with an exponential function

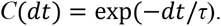

### Cell death quantification

FITC blobs in follicle were identified by characteristics such as high FITC intensity, spherical morphology and similar in size to GC. FITC blob and GC in FITC stained follicles were segmented using labkit in FIJI and manually checked to ensure consistency. The result was sent to Imaris to compute the count in 3D. A ratio between FITC blob count to GC count was taken to get the number of FITC blob per follicle. For co-localisation study with cCASP3, follicles stained and imaged with FITC were embedded with 2% agarose in the microwell before they were fixed and stained with cCASP3 and imaged in the same orientation in the microwell. The two images were co-registered before identifying the corresponding FITC blob and cCASP3 pair. In SEM, GC cell death in follicle was identified by characteristics such as nuclear condensation, shrunken size, dark pigmentation, blebbing and loss of structural integrity in organelles or cytoplasm. They were manually counted, along with normal GC with visible nucleus, and was taken as a ratio to get cell death per follicle.

### GC doublet assay

GCs were isolated from follicles by rupturing follicles in isolated ovaries using a 27-gauge needle. GCs released into the isolation buffer were filtered using a 30 µm cell strainer (Miltenyi Biotec, 130-098-458) before they were centrifuged at 300 g for 5 min. Isolation buffer was aspirated and the GC cell pellet was resuspended with 1 mL of growth medium. A 1:20 dilution was carried out using the GC suspension and seeded into an agarose microwell. GC doublets were located on the IX83 widefield microscope (Olympus) with a UPLSAPO S2 60x objective fitted with a DP23M camera (Olympus). Drugs were added following acquisition of timepoint 0 min. The GC doublets were tracked and imaged at 1 µm z step for 11 µm in depth every 30 min for 90 min.

For each GC in the doublet across time, a circle was drawn at the focal plane of the GC using the oval selection tool in FIJI to extract the radius and xy coordinates of its centre. The z slice was also recorded. The distance between the two GC in the doublet, *D*, was calculated by taking the distance between the two centres of the GC. Assuming the contact is small, and two cells are perfect balls overlapping by just minimal portion, then we could derive their contact angle, *θ* as

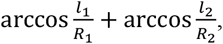

where *R*_1_, *R*_2_ are the radius of two cell balls and

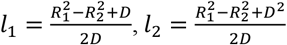

are the distances from the interface surface to the two cell centres, respectively. The change in *D* and *θ*between 0 min and 90 min was calculated and plotted for control and ADH-1.

### Follicle growth analysis

Isolated follicles cultured in 3D were imaged every day on the EVOS™ M5000 Imaging System (Thermofisher) using the 10X objective for up to 7 days. Microtissues were imaged on the SMZ1000 stereoscope (Nikon) with a Plan Apo 1X objective on 4X zoom fitted with a TrueChrome 4K Pro camera (Tucsen). Follicle diameter, D, was measured by taking an average of the major and minor axis in the drawn polygonal selection of the follicle on FIJI. Growth rate, *γ*, of the follicle was defined as the change of diameter, d*D*, over a period of time, dt, normalized by the diameter: γ = *dD*(*t*)/*D*(*t*)*dt*, which is equivalent to *d* ln*D*(*t*)/*dt* when dt is small. In practice, we calculated the growth rate of a follicle with diameter D at time t from the discrete time evolution as (ln(*D*(*t*)) − ln (*D* (*t* − Δ*T*)) /Δ*T*, where Δ*T* is the time interval between two consecutive time points and Δ*T* is usually 1 day.

### Lumen growth analysis

As the largest IG volume or lumen fraction grows with time non-linearly and we took logarithmic scale of both the quantity *Q* of interest and time *t* and used linear fitting to find the power *g* so that log*Q* = *g*log*t* + *k*. In the experiment (Fig.1 g-h), the growth power is fitted from 100 min after the initiation of timelapse. In the simulation, the growth power is fitted from the time point where the steady state is reached (Fig. 5g).

### Statistical methods

All means or medians from pooled data are plotted with error bars indicating SEM. Test of difference between two populations is one-sided Mann-Whitney U test, unless otherwise stated. Test of difference for drug treatment (before and after), or between middle and basal GCs is one-sided Wilcoxon signed rank test. In all tests, n.s. indicates non-significant, for p-value > 0.05.

## SUPPLEMENTARY INFORMATION

### Basic settings of the theoretical model

In this paper, since granulosa cells (GCs) exhibit fluid-like behaviour at the scale of follicle development, we model the follicle including luminal fluid as a binary fluid mixture, where one fluid component corresponds to the GC region and the other to the lumen. To describe this mathematically, we use the on-lattice spin representation, i.e., kinetic Ising model. Each lattice site *i* is assigned a spin *s*_*i*_ of either +1 or −1, representing the GC and lumen components, respectively. To account for the presence of the oocyte surface and follicle periphery, we assume a finite-size system with a fixed boundary mimicking their shapes (see Fig. SI1a). In this description, the fraction of lumen components (lumen fraction) is represented by the total number of negative spins per lattice site, *i*.*e*., *f*_*L*_ = (1/*N*) ∑_*i*_(−*s*_*i*_ + 1)/2 (summation runs over *i*) with the total number of sites *N*. In what follows, we focus on a regular square lattice, but we expect that the key results such as power-law behaviour do not depend on the choice of lattice type.

The dynamics is implemented via a kinetic Ising model that conserves the lumen fraction *f*_*L*_. Specifically, we use the Kawasaki dynamics, where the system evolves through local exchanges of nearest-neighbour spin pairs, minimizing the Hamiltonian determined by spin– spin interaction energy and effective temperature. The interaction energy, denoted as *J* , corresponds to the interfacial tension and reflects adhesion between neighbouring GCs. The effective temperature, denoted as *T*, reflects intrinsic stochastic motility of cells. For example, a large *J* indicates strong cell-cell adhesion, suppressing mixing between cells and luminal fluid and promoting phase separation, whereas a high *T* reflects strong random motility of cells, allowing cells to overcome cell-cell adhesion and enhancing mixing between cells and luminal fluid. Further details of this Hamiltonian are described in the following section. Based on our experimental data (Fig. 4f–h), we assume the existence of a spatial gradient in *J*/*T*, with the lowest values near the oocyte and the highest at the basal GCs. This gradient is incorporated in our simulation (Fig. 5a, left), as described further in the following sections. For the initial state, spin values are randomly assigned to lattice sites, with the lumen fraction *f*_*L*_ set below 0.1, in accordance with our quantification at population level (Fig. 1c).

### Details of the model and numerical simulations

The Hamiltonian of the system is given by

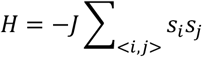

where < *i, j* > denotes nearest-neighbour pairs and *J* is the coupling strength between spins. Kawasaki dynamics is implemented as the exchange of the spin values between the nearest neighbours, with probability

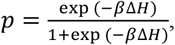

where Δ*H* is the change in free energy upon spin swapping, *β*= *J*/(*k*_*B*_*T*) (here, *k*_*B*_ represents the Boltzmann constant and, in what follows, we use the unit system with which *k*_*B*_ = 1) is the ratio of the coupling strength *J* (between like components) to the effective temperature *T*, indicating the effective intercellular tension that separates cell and lumen compartments. The value of *β* is prescribed to each site with a radial gradient:

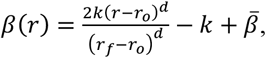

where and *d* is the lattice dimension, *r*_*f*_ and *r*_*o*_ are the radii of the simulated follicle and oocyte, respectively, *r* is its distance to the follicle center, 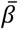 is the global baseline value of *β* , and *k* controls the steepness of the spatial gradient of *J*/*T*. Experimentally, *J* corresponds to the combined effect of GC junctional length and N-cad continuity (Fig. 4f, g), while *T* corresponds to the instantaneous velocity of GCs (Fig. 4h).

**Fig. SI1.**
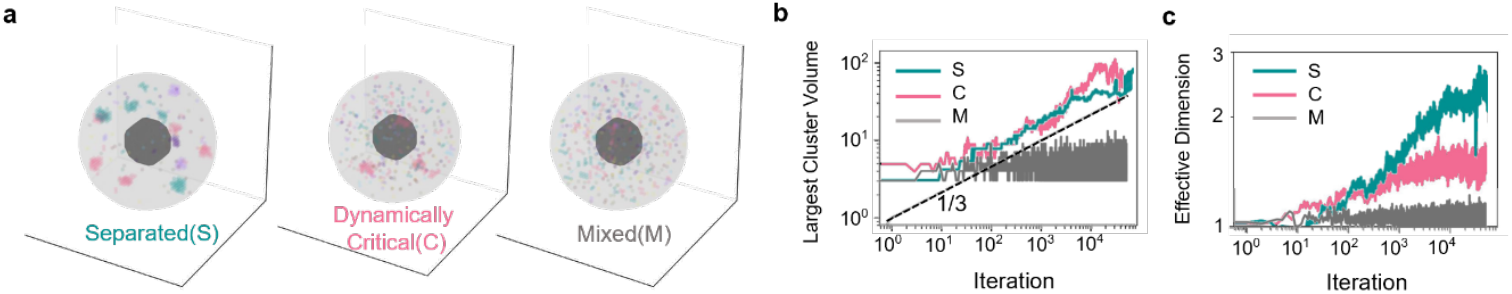
a, Representative snapshots of simulation reaching three steady states, separated (S), dynamically critical (C) and Mixed (M) on 3D lattice. **b-c**, Plots of temporal evolution of largest cluster volume (**b**) and effective dimension (**c**) in the three steady states.

Simulations are initialized by randomly assigning *s*_*i*_ = −1 (lumen component) on lattice sites with a probability equal to the given lumen fraction *f*_*L*_ (either the constant or initial) and *s*_*i*_ = 1 (cell component) with a probability 1 − *f*_*L*_. At each iteration of timestep, all pair of nearest neighbours are swapped with a probability *p* in a sequence randomly determined. The follicle diameter is set to 96 in 2D and 80 in 3D, while the oocyte diameter is set to 3/8 of the follicle diameter. From comparing the correlation time of the system at critical boundary (Fig. 5h) with the correlation time found in experimental near-critical regime (Fig. 2b), we could roughly estimate the timescale of each iteration corresponds to 10 seconds in real time.

### Results of Kawasaki dynamics in two or three dimensions with constant lumen fraction

First, we investigated the case where the lumen fraction was kept constant. The results on 2D and 3D lattice are qualitatively similar, resulting in three distinct steady states: namely, separated (S), dynamically critical (C), and mixed (M). Representative 2D lumen configurations of these states are shown in Figs. 5a and S1a and Supplementary Videos 10 and 11. The separated state is characterised by the nucleation of stable lumen clusters across the system, exhibiting a clear phase separation between cell and lumen components. The mixed state is the mixture of the two components, characterised by transient fluctuations in lumen dynamics without the formation of stable lumen clusters. The dynamically critical state represents the intermediate condition between phase separation and mixing, characterised by the formation of large morphologically unstable lumen clusters near the region of low *J*/*T* (*i*.*e*., near oocyte). In addition, in the dynamically critical state, small lumens still persist.

Regarding the dynamical features, in both the separated and dynamically critical states, the size of largest cluster grows over time approximately as a power-law with the exponent 1/3 (Figs. 5b and SI1b for 2D and 3D, respectively). However, only in separated state, the effective dimension of lumen clusters continuously increase toward the spatial dimension of the system (Figs. 5c and SI1c for 2D and 3D, respectively), indicating that lumen cluster boundaries are progressively rearranged into a more compact, spherical shapes; in contrast, in the dynamically-critical and mixed states, the effective dimension of lumen remains low, indicating that the shape of lumen remains complex due to the rapid rearrangement of lumen boundary.

We found that the global baseline value 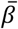 of *β*, the steepness *k* of the spatial gradient of *β*, and the lumen fraction *f*_*L*_ collectively determine which of the aforementioned steady states emerges. The results are summarized in a three-axes phase diagram (Fig. 5e). The phase boundary between the dynamically critical state and the other two states was identified by a sharp improvement in the goodness-of-fit to a power-law for the complementary cumulative distribution function (cCDF) (Fig. 5f). Notably, the dynamically critical state emerges only when *k* is nonzero.

From 2D projections of the phase diagram (Fig. 5f), we found that the dynamically critical state lies between the mixed and separated states, and that its domain expands with increasing steepness *k*. A reduction in the global value 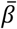 correspondingly enlarges the mixed state region, as expected when fluids are highly dynamic or interfacial tension is low.

### Results of 2D Kawasaki dynamics with varying lumen fraction

On a 2D lattice, we explored the role of lumen fraction constantly varying over time. Biological processes such as GC death and proliferation could contribute to changes in lumen formation, as partially observed experimentally (Fig. 3j). We suppose cell death to be a major contributor to the increase in the lumen fraction. The role of cell proliferation is straightforwardly assumed to decrease the lumen fraction by shrinking the cluster boundary. We also analysed the special distribution of cell death and cell proliferation in isolated follicles stained with cCASPS3 and p-H3Ser28 respectively (Extended Data Fig. 9a-b) and found that in small follicles, cell death occurs mostly near the oocyte, with an exponential tail in its radial distribution profile (Extended Data Fig. 9c), while cell proliferation is more uniformly distributed in space (Extended Data Fig. 9d).

Cell death is incorporated as lumen creation (with an exponential radial distribution) whereas cell proliferation is incorporated as lumen elimination (with a uniform spatial distribution). The number of cell components (*s*_*i*_ = 1) flipping to lumen components (*s*_*i*_ = −1) per timestep, denoted as *n*_*c*_, is the lumen creation rate and the number of lumen components (*s*_*i*_ = −1) flipping to cell components (*s*_*i*_ = 1) per timestep is the lumen elimination rate. The ratio of the elimination to creation rates is referred to the elimination/creation (E/C) ratio, ζ. The flipping probability *p*_*c*_ for lumen creation follows an exponential decay:

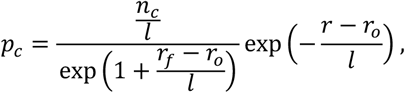

where *l* is the characteristic decay length. In our simulations, *l* is set to *r*_*o*_ /4. The flipping probability for lumen elimination is uniform across the space and given by ζ*n*_*c*_*/N*, where *N* is the number of cell components (*i*.*e*., the sites with *s*_*i*_ = 1) that has at least one lumen neighbour (*i*.*e*., the site with *s*_*i*_ = −1).

We performed the numerical simulations with a set of parameter values that reproduce the dynamically critical state under conditions where lumen creation and elimination are absent. To assess the impact of introducing these new processes, we systematically varied the lumen creation and elimination rates. The results are summarized in the phase diagram (Extended Data Fig. 9e) with two control parameters: lumen creation rate and the E/C ratio. E/C ratio =1 means lumen fraction is globally conserved while locally cell-lumen turnover occurs; E/C ratio <1 means that lumen creation is more than elimination, indicating an increase in lumen fraction over time, which is more relevant to the development stage of secondary follicle. We confirmed that if lumen creation is slow, the original steady states almost remain unaffected. If lumen creation is relatively fast, especially with a decreasing lumen fraction, the system evolves into mixed state. If lumen is created slowly with few eliminations of lumen, the system transitions to a new state, characterised by a super linear expansion of the largest cluster. The tricitical point for the super linear expansion to occur for E/C ratio ∼1 has a lumen creation rate about 0.01 per iteration, corresponding to 0.1 GC death ratio over 1 min which is trivial and not realistic. The results suggest that lumen growth in follicles observed in the experiments is primarily governed by the conservative fluid dynamics driven by cell-lumen interactions, and the non-conservative processes like cell death/proliferation should occur relatively slowly This enables the system to remain steady and progressively transition from the dynamically critical to separated states by the slow increase in lumen fraction.

## SUPPLEMENTARY VIDEOS

**Supplementary Video 1: Video of 3D rendered gap configurations for a secondary follicle (<180 µm)**.

**Supplementary Video 2: Timelapse video of a FITC-labelled secondary follicle**.

**Supplementary Video 3: Timelapse video of a FITC-labelled secondary follicle (<180 µm), showing dynamic IGs at short timescales**.

**Supplementary Video 4: Timelapse video of a FITC-labelled tertiary follicle (>180 µm), showing stable IGs at short timescales**.

**Supplementary Video 5: Timelapse video showing GC dynamics in a H2B-mCherry labelled secondary follicle**.

**Supplementary Video 6: Timelapse video captured by QPI, showing GC dynamics in a secondary follicle**.

**Supplementary Video 7: Timelapse video captured by QPI, showing cell rounding and changes in RI within a FITC-labelled tertiary follicle**.

**Supplementary Video 8: Timelapse video of a FITC-labelled tertiary follicle, showing FITC-blobs gradually merging with the IG**.

**Supplementary Video 9: Timelapse video captured by QPI, showing T1 transition in basal GCs of a follicle**.

**Supplementary Video 10: Timelapse video for simulation of IG growth leading to separated (S) state**

**Supplementary Video 11: Timelapse video for simulation of IG growth leading to a dynamically critical (C) state**

**Supplementary Video 12: Timelapse video of a FITC-labelled secondary follicle treated with ADH-1**.

**Supplementary Video 13: Timelapse video captured by QPI, showing GC dynamics in an ADH-1-treated secondary follicle**.

## Notes

### Competing Interest Statement

The authors have declared no competing interest.

